# Getting a head: Evidence for a conserved anterior head patterning gene network in arthropods

**DOI:** 10.64898/2026.04.25.720801

**Authors:** Bradleigh M. J. Cocker, Andrew D. Peel

## Abstract

The head of chelicerates, such as spiders, scorpions and mites, is composed of the ocular, chelicerae and pedipalp segments and is considered to be homologous to the procephalon of insects which comprises the ocular, antennal and intercalary segments. Head segmentation in the spider, *Parasteatoda tepidariorum*, is a dynamic process in which a single stripe of expression of the *P. tepidariorum hedgehog* (*hh*) gene splits twice to form three separate stripes, which help pattern the three spider head segments. This dynamic *hh* stripe splitting process is dependent on spider homologues of the transcription factors *orthodenticle* (*otd*) and *odd-paired* (*opa*). Here we investigate the conservation of this dynamic patterning mechanism in two insect models: the hemimetabolous pea aphid, *Acyrthosiphon pisum*, and the holometabolous red flour beetle, *Tribolium castaneum.*. We show that insect *hh*, *otd* and *opa* homologues are expressed in a highly conserved temporal and spatial pattern during procephalon development in these insects. Our data are consistent with an ancestral insect state in which a single *hh* stripe splitting event underpins patterning of the ocular and antennal segments, followed by *de novo* formation of the intercalary *hh* stripe. Using parental RNAi in *T. castaneum*, we show that *hh*, *otd* and *opa* homologues exhibit striking similarities in their regulatory interactions during spider and insect head/procephalon segmentation. Our data suggest that *hh*, *otd* and *opa* homologues contribute to an ancient and largely conserved gene network controlling head/procephalon patterning in arthropods. We discuss the implications of these data for our understanding of the origin and evolution of the arthropod head, and propose a new model for the evolution of anterior patterning in holometabolous insects.

## Introduction

Classically referred to as “The arthropod head problem” or the “Endless dispute”, our understanding of arthropod head evolution has progressed, but some key aspects remain unresolved (1–3). In particular, questions remain regarding the anterior of the Euarthropod head.

The Euarthropod head exists in two configurations: the three-segment head of Chelicerata (spiders, scorpions and mites) and the six-segment head of Mandibulata, which includes Myriapoda (centipedes and millipedes) and Pancrustacea (crustaceans and insects) (Fig.1). Disagreement remains over whether the most anterior ‘segment’, the ocular segment, should be considered a true segment, and whether an additional segment might exist anterior to this, giving rise to the labrum in insects. In this paper we’ll be assuming the hypothesis outlined in Figure 1. Under this hypothesis, conservation of *Hox* gene expression (4–7) as well as morphological evidence (8), suggests that the three segments of the chelicerate head are homologous to the anterior three segments of the mandibulates (Fig.1), which are collectively referred to as the procephalon (4–6, 9).

**Figure 1.**
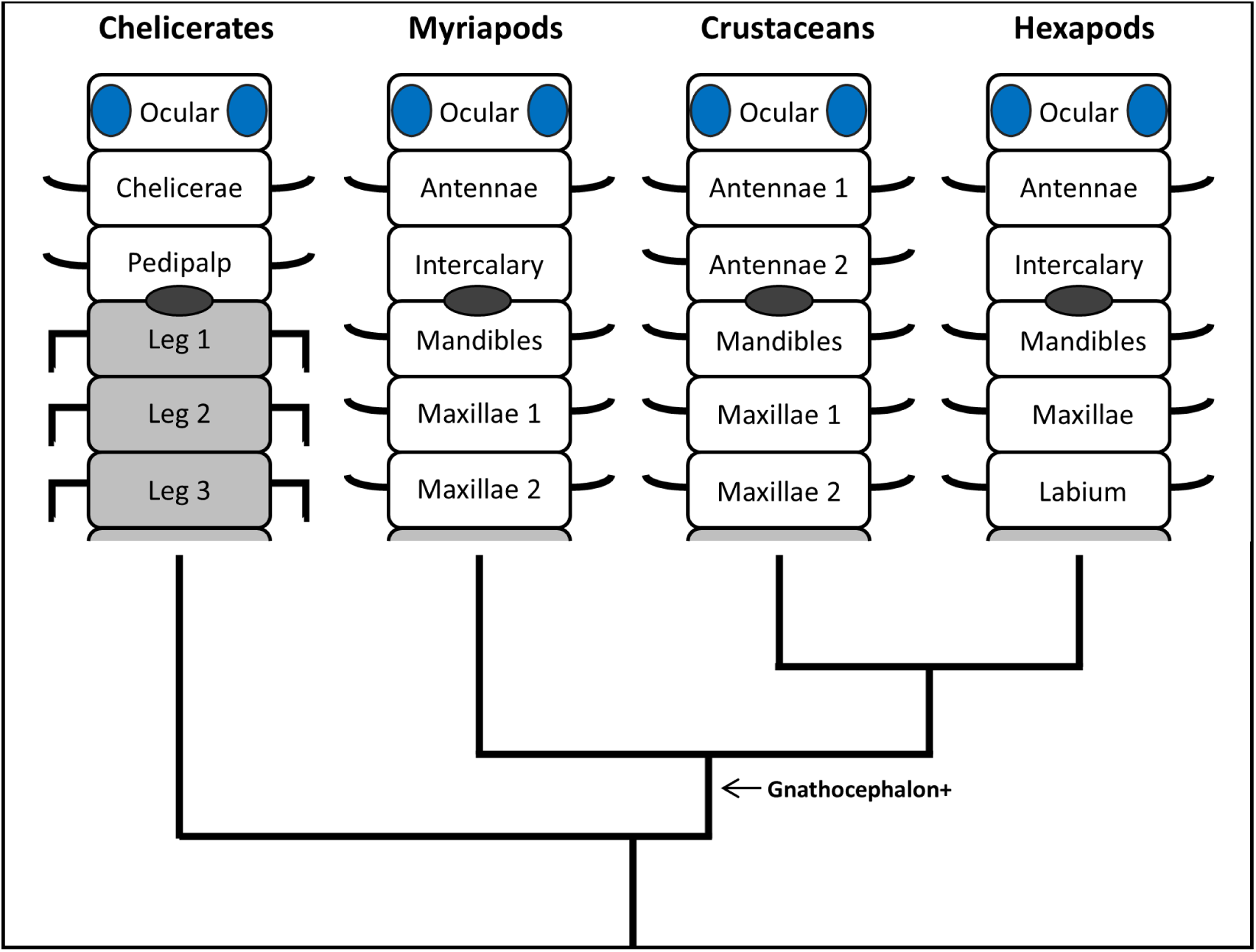
Homology of head segments across Euarthropoda. Diagram aligning the homologous segments of chelicerate, myriapod, crustacean and hexapod heads. Head segments are white, leg segments are shaded grey. The position of the mouth is indicated by a grey oval. Each appendage-bearing segment is marked with a simplified appendage. Phylogeny of each group is displayed according to the Pancrustacea hypothesis. The displayed phylogeny has been simplified to place insects besides crustaceans rather than within the paraphyletic crustacean clade to allow for alignment. An arrow marks the point at which the gnathocephalon is believed to have evolved.

The molecular mechanisms generating the head of chelicerates and the procephalon of mandibulates appear distinct to those patterning the posterior head segments in mandibulates (gnathal segments) and the arthropod trunk (for a recent review of trunk segmentation see Clark et al. (10)). The *hedgehog* signalling pathway appears to play a prominent role in head/procephalon development, in particular the pathway’s ligand encoded in arthropods by the gene *hedgehog* (*hh*). In the spider model *Parasteatoda tepidariorum (P. tepidariorum*), generation of the three chelicerate head segments depends on a dynamic “stripe splitting” mechanism involving the *P. tepidariorum* homologue of *hedgehog* (*Pt-hh*). Specifically, a single stripe of *Pt-hh* expression splits twice to produce three stripes of expression marking the posterior boundary of the three future *P. tepidariorum* head segments (11). In the myriapod centipede, *Strigamia maritima* (*S. maritima*), procephalon segments are patterned through a similar double *hh* stripe splitting process, albeit the segments are specified in a different order (12). In contrast, in all other arthropods studied only one *hh* stripe splitting event occurs, giving rise to the two most anterior head segments. This includes a second myriapod, the millipede *Glomeris marginata* (*G. marginata*) (13), numerous chelicerates including some spiders (14), and all insects studied to date (15, 16).

The comparative *hh* expression data outlined above imply two possible evolutionary scenarios: 1. Double *hh* stripe splitting might be ancestral to arthropods, with a reduction to a single stripe splitting event occurring convergently in multiple lineages. The reduction to a single stripe splitting event in insects and myriapods could be explained by the convergent reduction, and delayed development, of the third (“intercalary”) procephalon segment and loss of its antennal appendage (e.g. the second antenna of crustaceans) during the transition of these lineages to life on land (17). However, the variation in *hh* stripe splitting observed within terrestrial spiders/chelicerates is more difficult to explain under this scenario (14). Conservation of double *hh* stripe splitting would support the idea that procephalon segmentation is a process quite distinct to the patterning of trunk segments, consistent with these three segments having a unique evolutionary history; i.e., they could owe their origin to the serial duplication of a single anterior ancestral segment (5, 18). 2. In the alternative scenario, single *hh* stripe spitting is the ancestral state, with increases in the number of splitting events occurring independently in different myriapod and chelicerate lineages (14). Whichever scenario is true, it’s clear that *hh* stripe splitting is a conserved phenomenon across arthropods, at least in relation to patterning of the two most anterior head segments, and understanding its regulation and level of conservation should contribute to resolving the arthropod head problem.

Functional studies that offer insights into how *hh* stripe splitting is regulated are limited, with most information coming from a study on *P. tepidariorum* (11). In this spider, homologues of the genes *orthodenticle* (*Pt-otd*) and *odd-paired* (*Pt-opa*) function to maintain *Pt-hh* expression during stripe splitting and to promote *Pt-hh* stripe splitting respectively (11).

Published expression and functional data are consistent with a role for Orthodenticle and Odd-paired homologues in anterior and/or head development in insects (19–23). Orthodenticle (abbreviated to Otd in arthropods, Otx in vertebrates and called Oceilliless/Oc in *Drosophila*) is a homeobox transcription factor with a highly conserved role in patterning anterior body regions across animals (24, 25). Odd-paired’s vertebrate homologues are called Zinc finger of the cerebellum (Zic) proteins. Opa/Zic proteins are zinc finger transcription factors that are widely recognised to regulate hedgehog signalling (via direct binding to Gli proteins) and inhibit Wnt signalling (via direct binding to TCF family members) in various developmental contexts, including in vertebrate brain development (26). Recently, Odd-paired has also been shown to act as a temporal ‘pioneer’ factor in insects, acting globally to open up chromatin and change the accessibility of a large subset of enhancers to other transcription factors, thus facilitating global changes in gene regulatory networks (27, 28). Odd-paired appears to do this in collaboration with, or downstream of, Zelda, during early embryogenesis along both the anterior-posterior and dorsal-ventral axis (27). A lot of attention has focused on Odd-paired’s global temporal role in the process of ‘frequency doubling’ during trunk segmentation (20, 28–30). Intriguingly, and perhaps of more relevance to this study, two recent studies suggest that Odd-paired may have an earlier role acting as a pioneer factor specifically in the anterior, opening early acting anterior and head-related enhancers for regulation (21, 31).

To determine whether the head gene network operating in *P. tepidoriorum* might be conserved across the Euarthropoda, we investigated the expression and function of homologues of *hh*, *otd* and *opa* in insects. The classical insect model *Drosophila melanogaster* (*D. melanogaster*) displays a derived larval head which is involuted and highly reduced (32). This evolutionary adaptation complicates and limits the value of *D. melanogaster* studies in regards to understanding the origin and evolution of insect head segmentation. Therefore, we studied head segmentation in two alternative insect models; the holometabolous red flour beetle *Tribolium castaneum* (*T. castaneum*) and the hemimetabolous pea aphid *Acyrthosiphon pisum* (*A. pisum*).

In comparison to *D. melanogaster*, *T. castaneum* larvae develop a fully everted head with segments and appendages morphologically similar to adults (32). Furthermore, functional analysis of head segmentation genes is possible through RNA interference (RNAi) (22, 23, 32–35). RNAi against *T. castaneum* homologues of each chelicerate head segmentation gene - *Tc-hedgehog* (*Tc-hh*), *Tc-orthodenticle-1* (*Tc-otd1*) and *Tc-odd-paired* (*Tc-opa*) - cause head reduction phenotypes, specifically deleting procephalic structures in the case of *Tc-hh* and *Tc-opa* RNAi (20, 22, 35). In addition, expression of each gene correlates with the developing procephalon (20, 22, 35, 36). However, so far these genes have been studied separately, and the relative expression patterns and functional interactions between these genes in the head have not been examined in any detail.

In contrast to *T. castaneum*, *A. pisum* is a hemimetabolous insect, meaning that it undergoes partial metamorphosis via a sequence of nymphal stages with no pupal stage (37). Furthermore, *A. pisum* can reproduce through both asexual viviparous development and sexual oviparous development (38, 39). *A. pisum* asexual development is an evolutionarily novelty and appears to utilise a distinct developmental program to the ancestral sexual morph, with differences in the expression of early developmental genes such as the *A. pisum* homologues of *caudal*, *hunchback* and *orthodenticle* (*Ap-otd*) (39, 40). Therefore, studying the chelicerate head segmentation network in *T. castaneum* and the asexual mode of *A. pisum* development allows the conservation of the chelicerate head segmentation network to be assessed and compared between two highly divergent insect models.

We find that the chelicerate head segmentation network gene homologues are broadly conserved at the level of gene expression patterns during anterior procephalon segmentation in both *T. castaneum* and *A. pisum*. Furthermore, we find that each gene has conserved functions in head segmentation between *T. castaneum* and *P. tepidariorum.* This is consistent with *hh*, *otd* and *opa* constituting an ancient and conserved gene network controlling head/procephalon segmentation in arthropods. This has important implications for our understanding of the origin and evolution of the arthropod head and the evolution of anterior-posterior patterning in insects.

## Results

### Expression of *Tc-hh* overlaps with *Tc-otd1* and *Tc-opa* throughout *T. castaneum* development

In order to investigate whether the *P. tepidariorum* head segmentation network is conserved in *T. castaneum*, we characterised expression of *T.castaneum* homologues of *hh*, (*Tc-hh*), *opa* (*Tc-opa*) and *otd* (*Tc-otd1*) throughout *T. castaneum* head development using multiplex HCR in situ hybridisation (20, 22, 35).

### The ocular and antennal *Tc-hh* stripes are produced from a single stripe splitting event

Firstly, we analysed the expression dynamics of *Tc-hh.* In agreement with previous work by Farzana and Brown (35), we observed that *Tc-hh* is first expressed from late blastoderm stages and appears as symmetrical stripes either side of the developing germ disc (Fig.2.Bv,Cv). The symmetrical stripes are localised in the ventral head field (future procephalon) to the anterior of the extending germband. We analysed the expression dynamics of this stripe across germband development using the number of trunk *Tc-hh* stripes as a readout for germband stage.

**Figure 2.**
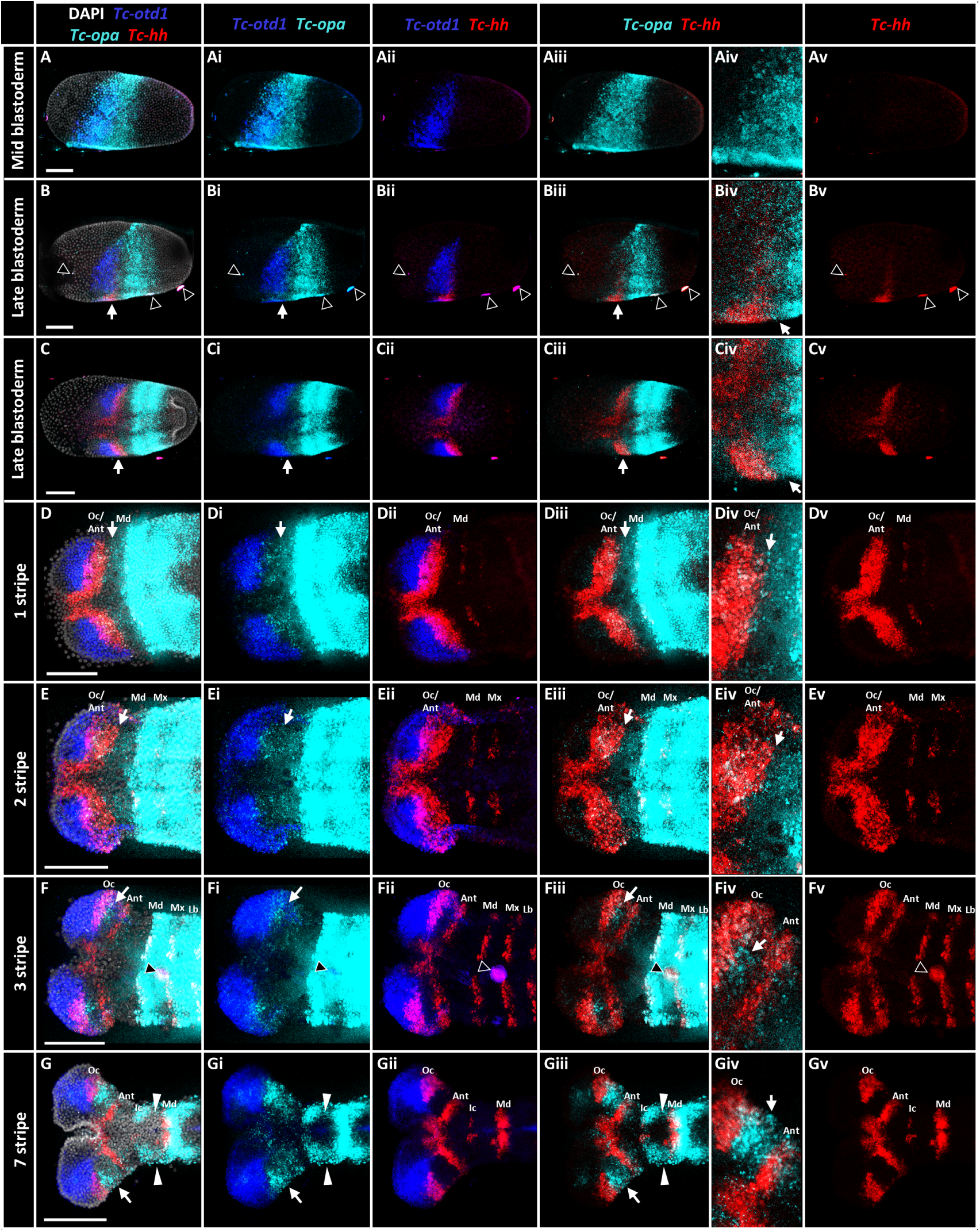
*Tc-hh* is expressed between overlapping domains of *Tc-otd1* and *Tc-opa* expression throughout *T. castaneum* procephalon segmentation. (A-Gv) Maximum projections of T. castaneum embryos of increasing age from youngest (A-Av) to oldest (G-Gv). Each embryo is stained for DAPI (grey) and expression of *Tc-otd1* (blue), *Tc-opa* (cyan) and *Tc-hh* (red). In germband stage embryos (D-Gv) Each *Tc-hh* stripe is labelled with its corresponding segment: Ocular (Oc), Antennal (Ant) Intercalary (Ic), Mandibular (Md), Maxillary (Mx) and Labial (Lb). Embryos are oriented with the anterior on the left. Embryos A and B are oriented with the dorsal side at the top and ventral side at the bottom, embryo C is oriented ventrally. White arrows mark the weak anterior domain of *Tc-opa* expression overlapping the ocular/antennal *Tc-hh* stripe. White triangles mark intercalary *Tc-opa* expression. Black triangles mark debris on the embryo (Bi-Bv, Fi-Fv). Scale bars: 100 µm.

*Tc-hh* remains expressed in the procephalon as a single stripe up to the appearance of the second sequential (maxillary) *Tc-hh* stripe (Fig.2.Dv,Ev.). Concurrently with the appearance of the third sequential (labial) *Tc-hh* stripe we found that the initial procephalic *Tc-hh* stripe splits to produce an anterior *Tc-hh* stripe marking the posterior boundary of the ocular segment and a posterior *Tc-hh* stripe marking the posterior boundary of the antennal segment (Fig.2.Fv). Therefore, the initial unsplit *Tc-hh* stripe was designated as the ocular/antennal stripe. No additional stripe splitting event was observed to generate the intercalary *Tc-hh* stripe. Instead, *Tc-hh* appears *de novo* in the intercalary segment around the time the seventh (1^st^ abdominal) *Tc-hh* stripe appears (Fig.2.Gv).

### The ocular/antennal *Tc-hh* stripe overlaps *Tc-otd1* and *Tc-opa* expression throughout head segmentation

Consistent with previous studies, we found that *Tc-opa* (20) and *Tc-otd1* (22) were expressed ubiquitously in early blastoderm stages (data not shown) and refined into wedge shaped domains by mid-blastoderm stages (Fig.2.Aii,Aiii). We found that the *Tc-otd1* expression domain overlaps with the anterior half of the *Tc-opa* expression domain (Fig.2.A-Ai). By late blastoderm development, the *Tc-otd1* expression domain splits medially into two distinct domains that mark the presumptive cephalic lobes (Fig.2.B-Bii, C-Cii). At this stage, it was previously shown that *Tc-opa* expression is lost from the anterior portion of the wedge and is confined to the gnathal segments (20). Due to the higher sensitivity of *in-situ* hybridisation chain reaction (HCR), we observed persistence of a previously unobserved weak *Tc-opa* expression domain in the presumptive procephalon, partially overlapping with expression of *Tc-otd1* (Fig.2.B,Bi, C,Ci, white arrow).

The ocular/antennal *Tc-hh* stripe appears between and overlapping expression of *Tc-otd1* to the anterior (Fig.2.Bii,Cii) and weak *Tc-opa* expression to the posterior (Fig.2.Biii,Biv,Ciii,Civ). This positional relationship persists up to the second sequential (maxillary) *Tc-hh* stripe stage with the ocular/antennal *Tc-hh* stripe partially overlapping both *Tc-otd1* and *Tc-opa* expression domains (Fig.2.Dii-Dv, Eii-Ev). During the third sequential (labial) *Tc-hh* stripe stage, the ocular *Tc-hh* stripe forms from cells in the anterior portion of the ocular/antennal stripe which co-express *Tc-otd1* (Fig.2.Eii,Fii,Gii). In contrast, the antennal stripe forms from cells at the posterior edge of the ocular/antennal stripe which overlap with the weak *Tc-opa* expression domain (Fig.2.Eiii,Fiii,Giii). *Tc-opa* is also expressed in the gap between the ocular and antennal segments and partially overlaps with the ocular *Tc-hh* stripe (Fig.2.Fiii,Fiv,Giii,Giv, white arrow). By the seventh (1^st^ abdominal) *Tc-hh* stripe stage, cells in the ocular *Tc-hh* stripe continue to express *Tc-otd1* (Fig.2.Gii) whilst *Tc-opa* is expressed in domains between each *Tc-hh* stripe (Fig.2.Giii) including in the intercalary and mandibular segments (Fig.2.Giii, white triangles).

### Knockdown of *Tc-hh, Tc-opa* and *Tc-otd1* expression using parental RNAi caused head reduction phenotypes

Next, we analysed the function of *Tc-hh*, *Tc-opa* and *Tc-otd1* by performing parental RNAi targeting each gene and targeting *gfp* as a control. As described in previous studies (20, 22, 35), we found that RNAi against each gene produced unhatched larval cuticles with head reduction phenotypes more frequently than cuticles produced from *gfp* RNAi (Fig.S.1.A,B,C). For *Tc-otd1* RNAi, we most frequently observed reduction of the procephalon and gnathocephalon (Fig.2.A) resulting in cuticles with no procephalon segments and loss or reduction of gnathocephalon segments (*Tc-otd3’*: N=109/166, 65.7%, *Tc-otd5’*: N=233/338, 68.9%) (Fig.S.1.Fiii). We observed the same phenotype less frequently in *Tc-opa3’* (N =4/89, 4.5%) and *Tc-hh* RNAi (N =3/219, 16.4%) cuticles (Fig.S.1.Eiii, Giii) but more frequently observed reduction or loss of the procephalon without reduction of the gnathocephalon (*Tc-opa 3’:* N =23/89, 25.8%, *Tc-opa5’*: N=5/22, 22.7%*, Tc-hh*: N=89/219, 40.6%) (Fig.S.1.B,C,Eii,Gii). Following analysis of cuticle phenotypes, we next analysed the effects of RNAi knockdowns of each gene on head gene expression and cellular morphology during embryogenesis.

### Knockdown of *Tc-opa* expression causes inhibition of *Tc-hh* stripe splitting and reduced head size

We first analysed the effects of *Tc-opa* RNAi on expression of *Tc-otd1* and *Tc-hh* during blastoderm development. We observed no difference in *Tc-otd1* expression between *gfp* control (Fig.3.A,B N = 10) and *Tc-opa* RNAi in blastoderm (Fig.3.C,D N = 12) and germband stage (not shown N = 12) embryos. In contrast, we observed a mild reduction in *Tc-hh* expression in *Tc-opa* RNAi blastoderm stage embryos (N = 10/12) when compared to *gfp* RNAi embryos of the same stage (N = 10 embryos). Specifically, the number of cells within the *Tc-hh* stripe, as well as the width (Fig.3.Aii, Cii) and length of the stripe (Fig.3.Bii, Dii), appeared reduced.

**Figure 3.**
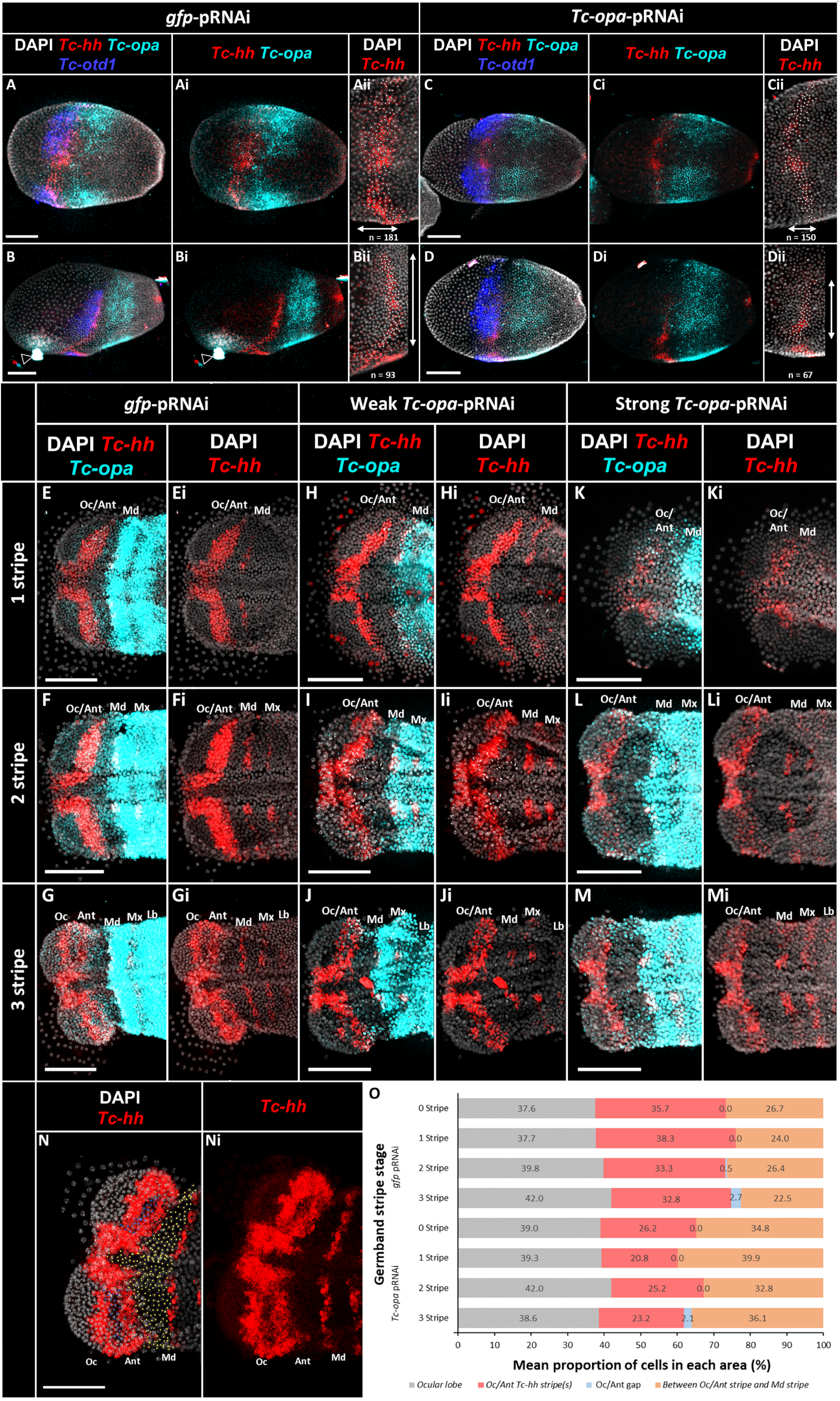
*Tc-opa* RNAi causes reduced ocular/antennal *Tc-hh* stripe expression, inhibits stripe splitting and reduces head lobe size. (A-Dii) Maximum projections of *T. castaneum gfp* RNAi (A-Bii) and *Tc-opa* RNAi (C-Dii) blastoderm embryos stained for DAPI (grey), *Tc-hh* (red), *Tc-opa* (cyan) and *Tc-otd1* (blue). In panels Aii-Dii, the approximate width (Aii, Cii) and length (Bii, Dii) of the *Tc-hh* stripe in each condition are indicated by double-sided arrows with each *Tc-hh* expressing cell labelled by a white dot. The number of *Tc-hh* positive cells in each stripe are displayed in each panel. (E-Mi) Representative maximum projections of *gfp-*RNAi (E-Gi) and *Tc-opa* RNAi (H-Mi) germband stage embryos aligned by stripe stage. *Tc-opa* RNAi embryos were observed with weak (H-Ji) and strong (K-Mi) head reduction phenotypes. Each *Tc-hh* stripe is labelled with its corresponding segment: Ocular (Oc), Antennal (Ant), Intercalary (Ic), Mandibular (Md), Maxillary (Mx) and Labial (Lb). (N, Ni) Representative *gfp*-RNAi germband procephalon used for cell counting with counted cells marked in each of the following categories: ocular lobe cells (grey), ocular/antennal *Tc-hh* cells (red), ocular/antennal gap cells (blue) and cells between the mandibular and ocular/antennal *Tc-hh* stripes (orange). (O) Graph of the mean proportion of cells in each area in *gfp*-RNAi and *Tc-opa* RNAi germband embryos from 0-3 stripe stages. Scale bars: 100 µm.

To understand the effects of *Tc-opa* RNAi on head segmentation, we next analysed expression of *Tc-hh* in germband embryos between the zero to three *Tc-hh* trunk stripe stages. We noticed three phenotypes in *Tc-opa* RNAi embryos: wildtype stripe splitting and no head lobe reduction (N=3/30, not shown), partially inhibited Oc/Ant stripe splitting with weak head lobe reduction (Fig.3.H-Ji, N=14/30) and inhibited Oc/Ant stripe splitting with strong head lobe reduction (Fig.3.K-Mi, N = 13/30). In contrast, no stripe splitting or head reduction phenotypes were observed in *gfp* RNAi embryos (Fig.3.E-Gi, N = 25). We further analysed the weak head reduction phenotype by quantifying the effect of *Tc-opa* RNAi on the mean proportion of cells in each area of the procephalon between the zero to three *Tc-hh* stripe stages (see methods and Fig.3.N-O). *Tc-opa* RNAi embryos displayed a significantly decreased mean proportion of ocular/antennal *Tc-hh* cells with respect to *gfp* RNAi control embryos (*t*(*3*) = 5.2, p = 0.014). In contrast, the mean proportion of cells between the mandibular and ocular/antennal *Tc-hh* stripes was significantly increased (*t*(3) = 4.9, p = 0.016). No significant difference was found between the proportions of ocular lobe cells (*t*(3) = 0.3, p = 0.751) or ocular/antennal gap cells (*t*(3) = 1.7, p = 0.184).

These findings suggest that *Tc-opa* is required for expression of *Tc-hh* along the posterior edge of the ocular/antennal *Tc-hh* stripe as well as ocular/antennal stripe splitting. In addition, *Tc-opa* appears to regulate head lobe size.

### Knockdown of *Tc-otd1* expression caused inhibition of *Tc-hh* stripe splitting and reduced head size

Previous studies found that *Tc-otd1* RNAi causes head reduction phenotypes (22, 23). However, it was shown that this was due to defects in dorsal-ventral axis specification leading to deletion of the ventral head field (23). To investigate whether head field deletion entirely explains *Tc-otd1* RNAi head reduction phenotypes, we analysed expression of *Tc-hh*, *Tc-opa, Tc-otd1* alongside *Tc-zerknüllt* (*Tc-zen)*, which is expressed in the serosa during blastoderm stage development (23), in *Tc-otd1* RNAi embryos. In *gfp* RNAi control blastoderm embryos, *Tc-zen* expression is visible throughout the serosa and illustrates the oblique dorsal-ventral boundary between the embryo and serosa (Fig.4.A-Aiii, arrowheads, N = 12/12). In contrast, we observed that *Tc-otd1* RNAi blastoderm embryos showed reduced expression of *Tc-zen* in the serosa and a shift in the embryo-serosa boundary such that the boundary appears less oblique (Fig.4. B-Biii, arrowheads, N= 13/15).

**Figure 4.**
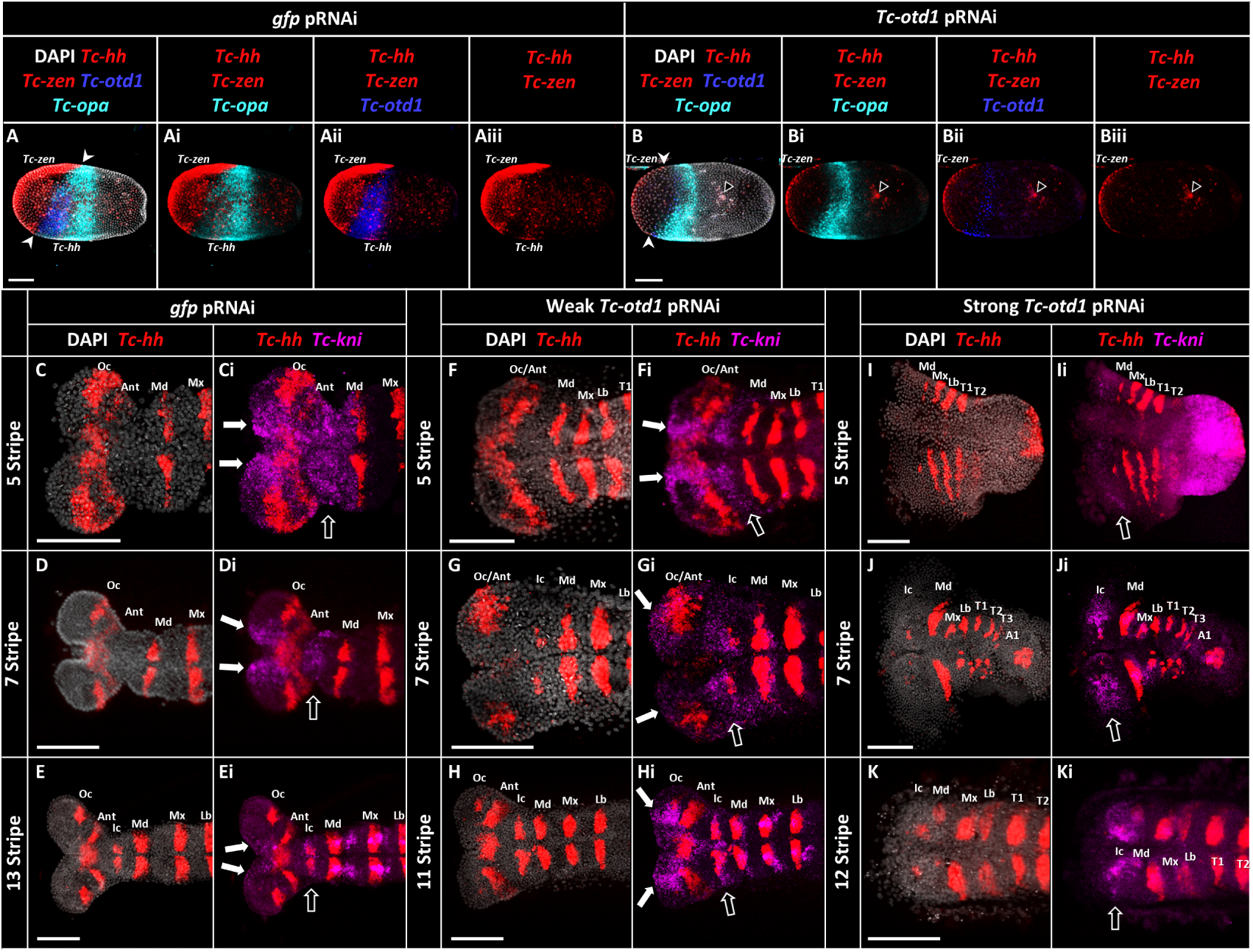
*Tc-otd1* RNAi causes head deletion and inhibited ocular/antennal stripe splitting independently. (A-Biii) Maximum projections of *T. castaneum gfp* RNAi (A-Aii) and *Tc-otd1* RNAi (B-Biii) blastoderm embryos stained for DAPI (grey), *Tc-hh* and *Tc-zen* (red), *Tc-opa* (cyan) and *Tc-otd1* (blue). Expression domains of *Tc-hh* and *Tc-zen* are labelled in each embryo. (C-Ki) Maximum projections of *T. castaneum* germbands stained for DAPI (grey), *Tc-hh* (red) and *Tc-kni* (magenta). *gfp-*RNAi embryos (C-Ei) of increasing stripe stage are aligned against *Tc-otd1* RNAi embryos with weak (F-Hi) and strong head reduction phenotypes (I-Ki) of similar age. Each *Tc- hh* stripe is labelled with its corresponding segment: Ocular (Oc), Antennal (Ant) Intercalary (Ic), Mandibular (Md), Maxillary (Mx) and Labial (Lb), Thoracic 1-3 (T1-T3) and Abdominal 1 (A1). All embryos are oriented with anterior to the left. White arrowheads mark the embryo-serosa boundary. White arrows mark anterior *Tc-kni* expression, Black arrows mark intercalary *Tc-kni* expression. Black triangles mark debris. Scale bars: 100 µm.

We next analysed the effects of *Tc-otd1* RNAi in germband stage embryos. In order to assess the extent of head deletion we analysed expression of *Tc-hh* and *Tc-opa* along with *Tc-knirps* (*Tc-kni*) which is expressed in two domains in the germband head in a stage-specific manner (41). We observed that expression of *Tc-kni* in *gfp* RNAi germband embryos appeared similar to wildtype embryos. Specifically, we observed *Tc-kni* expression in anterior domains in the medial region of each head lobe (Fig.4.Ci-Ei, white arrows) as well as a posterior domain in the intercalary and mandibular segments (Fig.4.Ci-Ei, black arrow). We compared the presence of these two *Tc-kni* expression domains in *gfp* RNAi and *Tc-otd1* RNAi embryos of similar stages to assess the severity of head deletion.

We identified two categories of *Tc-otd1* RNAi embryos: embryos with both *Tc-kni* expression domains present (weak *Tc-otd1* RNAi, N=10/19) and embryos with one or both *Tc-kni* expression domains absent (strong *Tc-otd1* RNAi, N= 9/19). When compared to *gfp* RNAi embryos of similar stage, weak *Tc-otd1* RNAi germband embryos were observed with inhibited ocular/antennal *Tc-hh* stripe spitting (Fig.4.G, N=7/19), stripe splitting with reduced ocular *Tc-hh* stripe expression (Fig.4.F,H, N=2/19) or no head segmentation phenotype (not shown, N=1/19). In strong *Tc-otd1* RNAi embryos, which expressed some portion of the intercalary/mandibular *Tc-kni* expression domain (Fig.4.I-Ki), identity of each subsequent segment was assigned based on their position along the anterior-posterior axis. In addition to complete deletion of the head lobes, these embryos displayed disrupted morphology and *Tc-hh* stripe expression in segments of the gnathocephalon, trunk and abdomen (Fig.4.I-K). No head segmentation phenotypes were observed in *gfp* RNAi embryos (N=15/15).

Taken together, our findings support the previous finding that *Tc-otd1* RNAi causes head reduction phenotypes via deletion of the ventral head field. However, our results from carefully observing weak RNAi knockdowns suggest that this early d-v patterning role often obscures a later role and *Tc-otd1* RNAi also causes head reduction phenotypes through inhibition of ocular/antennal *Tc-hh* stripe splitting; this effect is most pronounced on the anterior/ocular side of the stripe where *Tc-otd1* is expressed. Therefore, we conclude that *Tc-otd1* functions to preserve the ventral head field, promote ocular/antennal stripe splitting and maintain *Tc-hh* expression in the ocular stripe.

### Knockdown of *Tc-hh* expression caused inhibition of ocular/antennal *Tc-hh* stripe splitting and *Tc-opa* expression

We next investigated the effect of *Tc-hh* RNAi on ocular/antennal *Tc-hh* stripe splitting as well as expression of *Tc-opa* and *Tc-otd1*. In blastoderm stage embryos we did not observe any differences in *Tc-opa* or *Tc-otd1* expression between *gfp* RNAi (Fig.5.A-Aii, N=7) and *Tc-hh* RNAi (Fig.5.B-Bii, N=5) embryos, suggesting that *Tc-hh* does not regulate expression of either gene at this stage.

**Figure 5.**
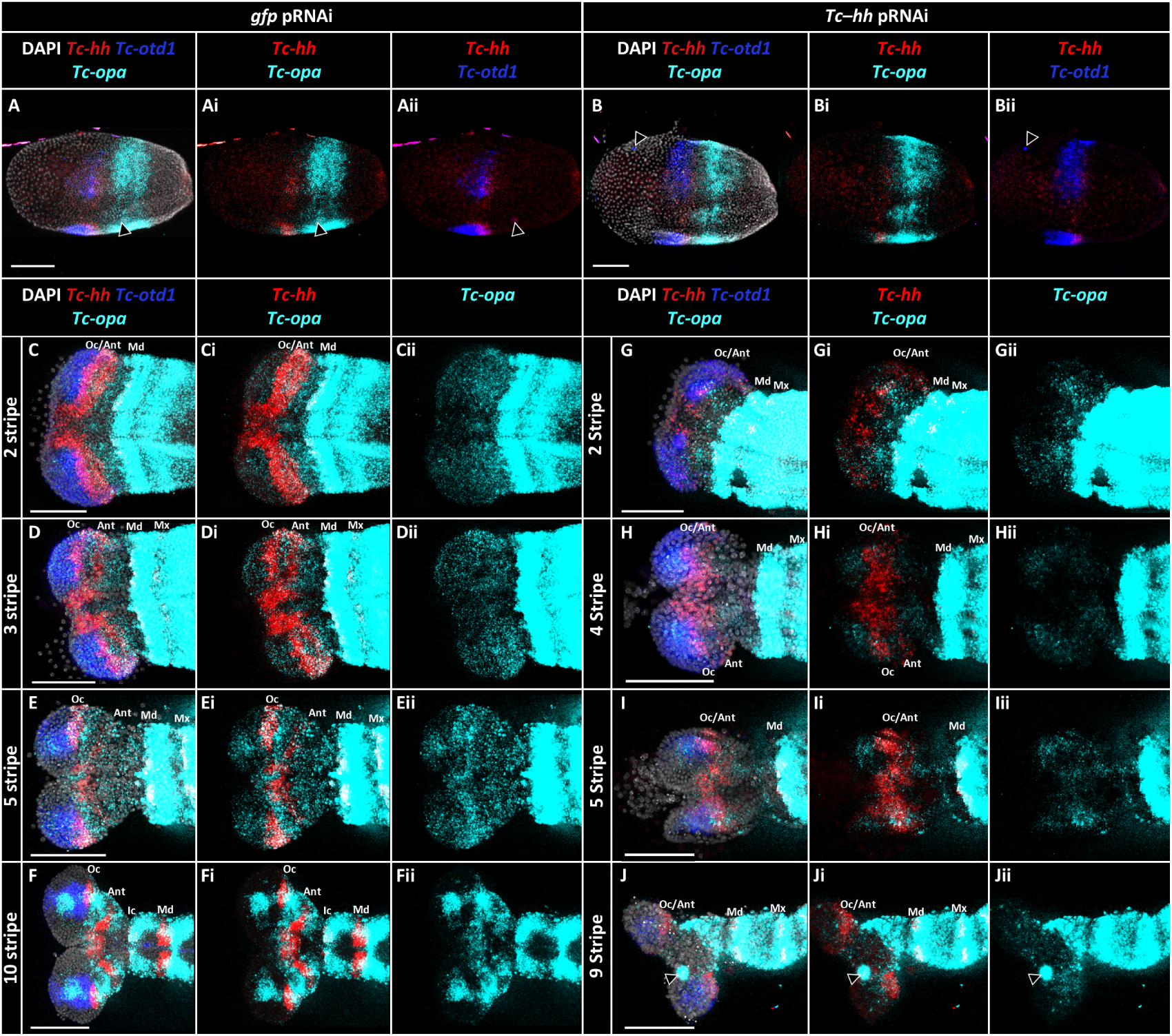
*Tc-hh* RNAi inhibits ocular/antennal stripe splitting and reduces ocular/antennal *Tc-opa* expression. (A-Bii) Maximum projections of *T. castaneum gfp* RNAi (A-Aii) and *Tc-otd1* RNAi (B-Bii) blastoderm embryos stained for DAPI (grey), *Tc-hh* (red), *Tc-opa* (cyan) and *Tc-otd1* (blue). (C-Jii) Maximum projections of *T. castaneum gfp* RNAi embryos (C-Fii) of increasing stripe stage aligned against *Tc-hh* RNAi embryos of similar age (G-Jii). Each *Tc-hh* stripe is labelled with its corresponding segment: Ocular (Oc), Antennal (Ant) Intercalary (Ic), Mandibular (Md), Maxillary (Mx) and Labial (Lb), Thoracic 1-3 (T1-T3) and Abdominal 1 (A1). All embryos are oriented with anterior to the left. Black triangles mark debris. Scale bars: 100 µm.

In germband stage embryos, we observed that *Tc-hh* expression appeared reduced but not absent in *Tc-hh* RNAi embryos (Fig.5.Gi-Ji) compared to *gfp* RNAi embryos (Fig.5.Ci-Fi) suggesting a relatively weak knockdown of *Tc-hh* expression. Despite this, we frequently observed that *Tc-hh* RNAi embryos displayed inhibited ocular/antennal stripe splitting, with embryos beyond the third (labial) *Tc-hh* stripe stage displaying poorly split or unsplit ocular/antennal stripes (Fig.5.Gi-Ji, N= 12/14). Normal ocular/antennal stripe splitting was observed in the remaining embryos (not shown, 2/14) and in all *gfp* RNAi embryos (Fig.5.Ci-Fi, N=16/16).

*Tc-opa* expression is visible in the procephalon at all stages in *Tc-hh* RNAi embryos with inhibited stripe splitting but appears reduced (Fig.5.Gii-Jii, N = 12/14) when compared to *gfp* RNAi embryos (Fig.5.Cii-Fii, N= 16/16). Specifically, *Tc-opa* expression appears reduced between the mandibular and antennal *Tc-hh* stripes and anterior to ocular *Tc-hh* expression. In contrast to *Tc-opa,* expression of *Tc-otd1* appeared relatively unchanged between *gfp* RNAi (Fig.5.C-F) and *Tc-hh* RNAi embryos (Fig.5.G-J) with minor differences in the size and shape of *Tc-otd1* expression domains.

Our findings suggest that *Tc-hh* promotes head segmentation through stripe splitting and may upregulate expression of *Tc-opa* in the ocular and antennal segments in germband embryos.

### The ocular and antennal segments are patterned via *Ap-hh* stripe splitting in asexual *A. pisum* development

We next characterised head segmentation in the asexual morphs of the hemimetabolous insect model *A. pisum*. (39)

To determine whether *hh* stripe splitting is conserved in asexual *A. pisum* head development, we analysed expression of *Ap-hh* and *Ap-Deformed* (*Ap-Dfd*), a homologue of the hox gene *Deformed* which marks the mandibular and maxillary segments in *T. castaneum* and *D. melanogaster* (42, 43). Asexual embryogenesis in *A. pisum* involves a complex series of movements regarding the position and morphology of the presumptive head (Fig.S.2.D-I, Ce). We staged each embryo by comparison with the detailed descriptions of Miura et al. (38).

We found that *Ap-hh* expression is visible around stage 7/8 and appeared as a stripe close to the anterior pole of the embryo (Fig.6.A-Aiii) as well as in a posterior expression domain (Fig.6.Aii, asterisk). In stage 10 embryos, we observed that the anterior *Ap-hh* stripe appeared to increase in size (Fig.6.B-Biii) and started to split by losing *Ap-hh* expression from cells in the posterior half of the stripe (Fig.6.Biii, blue). In stage 11 embryos, we observed that the anterior *Ap-hh* stripe had split into 2 distinct stripes (Fig.6.C-Ciii) which remained separate up to stage 16 and gave rise to the ocular and antennal segments (Fig.6.D-Diii). We did not observe any additional stripe splitting events for either stripe. Therefore, we assigned the initial anterior *Ap-hh* stripe as the ocular/antennal stripe.

**Figure 6.**
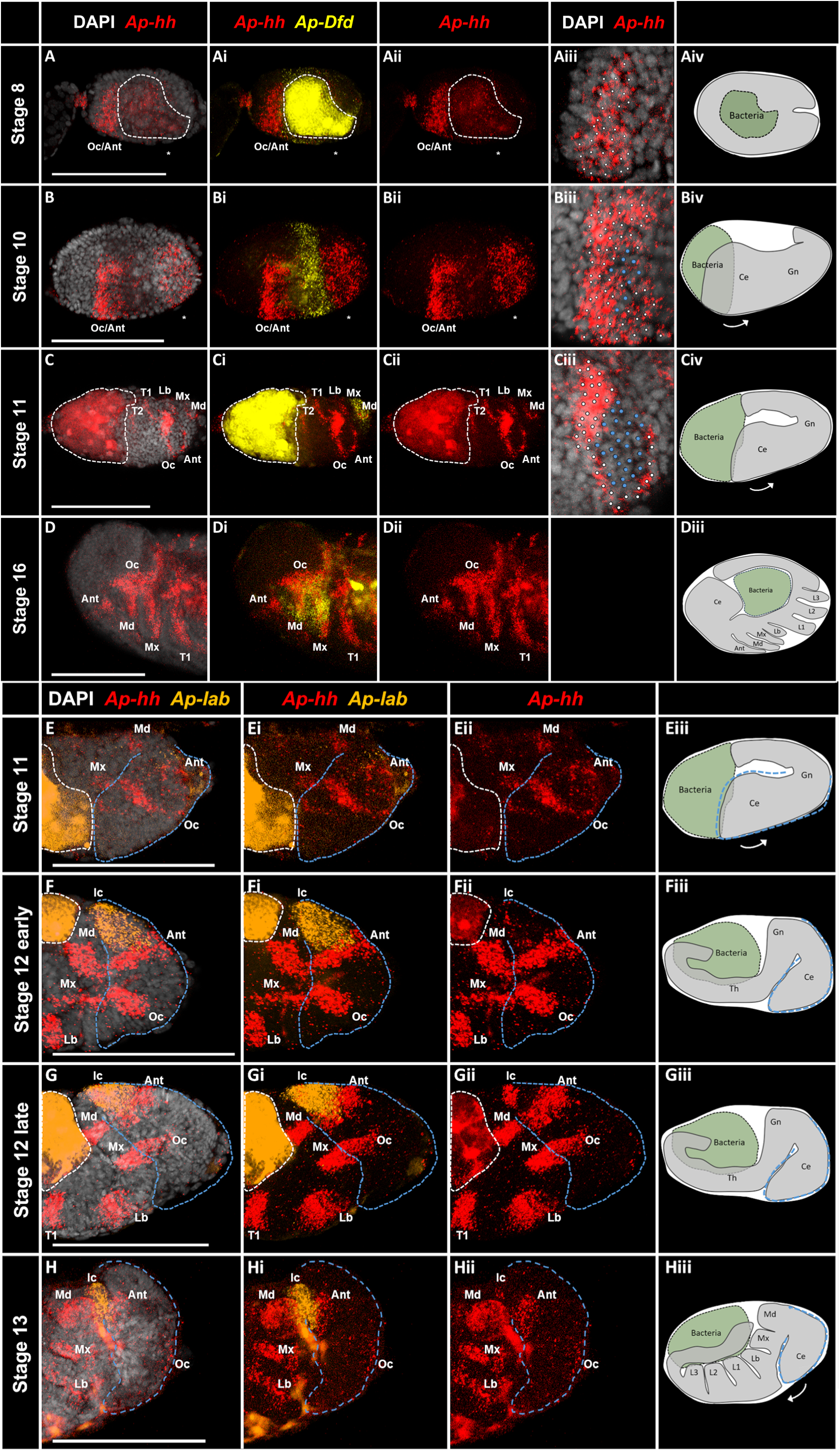
Conservation of ocular/antennal *Ap-hh* stripe splitting and *de novo* intercalary *Ap-hh* expression in viviparous *A. pisum* head segmentation. (A-Hii) Maximum projections of viviparous *A. pisum* embryos of increasing age stained for DAPI (grey), *Ap-hh* (red) and *Ap-Dfd* (A-Di, yellow) or *Ap-lab* (E-Hi, orange). (Aiii-Ciii) Cells spanning the ocular/antennal *Ap-hh* are marked as *Ap-hh*+ (white) or *Ap-hh*-(blue). Morphological diagrams (Aiv-Civ, Diii-Hiii) of corresponding stages are displayed alongside each row to aid interpretation. Each *Ap-hh* stripe is labelled with its corresponding segment: Ocular (Oc), Antennal (Ant) Intercalary (Ic), Mandibular (Md), Maxillary (Mx) and Labial (Lb), Thoracic 1-2 (T1-T2). All embryos are oriented with the anterior to the left. White dashed lines mark the position of the bacteriocyte where visible. Blue dashed lines indicate the position of the procephalon (E-H). White asterisk (*) marks posterior *Ap-hh* expression. Scale bars: 100 µm.

For *Ap-Dfd*, we observed a broad band of expression posterior to the Oc/Ant *Ap-hh* stripe from stage 8 onwards (Fig.6.Ai,Bi). In stage 11 embryos the *Ap-Dfd* expression domain overlaps two *Ap-hh* stripes that mark the first two sequential segments of the germband (Fig.6.C,Cii). We observed that between stage 11 and stage 16, *Ap-Dfd* remained specifically expressed in these two segments which began to develop appendages (Fig.6.D-Dii, Md, Mx). The position and morphology of these segments at stages 11 and 16 match the mandibular and maxillary segments previously identified by Miura et al. (38). Therefore, we assigned these *Ap-Dfd* expressing *Ap-hh* stripes as the mandibular and maxillary segments.

We next investigated whether the intercalary segment develops an *Ap-hh* stripe *de novo* as observed in *T. castaneum* (Fig.2.Gv) (44). We stained embryos for *Ap-hh* along with the *A.pisum* homologue of the hox gene *labial*, *Ap-lab*, which marks the intercalary segment in *D. melanogaster* and *T. castaneum* (44, 45). We found that *Ap-lab* expression first appeared between the end of stage 11 and the start of stage 12 as an expression domain between the antennal and mandibular *Ap-hh* stripes (Fig.6.Ei,Fi). In early stage 12 embryos, we observed the *de novo* appearance of *Ap-hh* expression at the posterior border of *Ap-lab* expression (Fig.6.Fi,Fii, Ic) which develops into a broad stripe later in stage 12 (Fig.6.Gii. Ic). Given the position of this segment between the antennal and mandibular segments, as well as the expression of *Ap-lab*, we assigned this *Ap-hh* domain as the intercalary stripe.

Taken together, these findings show that head segmentation is remarkably similar in *T. castaneum* and *A. pisum* as both involve an initial ocular/antennal *hh* stripe which splits a single time to produce separate ocular and antennal stripes, whereas the intercalary *hh* stripe appears later *de novo*.

### *Ap-otd* and *Ap-opa* are expressed overlapping *Ap-hh* during ocular/antennal stripe splitting

We next investigated the expression of *A. pisum* homologues of *otd* (*Ap-otd*) and *opa* (*Ap-opa*) in relation to *Ap-hh* expression.(11) Previous studies have shown that expression of *Ap-otd* is different between sexual and asexual development in *A. pisum*. Specifically, *Ap-otd* expression was shown to be absent until mid-late germband stages in asexual development (39, 40) which contrasts with the blastoderm expression of *Ap-otd* observed in sexual development of *A. pisum* (39) as well as the blastoderm expression of *Ap-otd* homologues in other arthropod models (11, 12, 22). Therefore, we reinvestigated whether *Ap-otd* was expressed during head segmentation in the divergent context of asexual *A. pisum* development using sensitive HCR in situ hybridisation.

In contrast to previous studies where *Ap-otd* expression was detected from stage 10/13 (39, 40), we found that *Ap-otd* was expressed as early as stage 7. In stage 7 embryos, *Ap-otd* expression appears in separate domains at the anterior (Fig.7.A,Aiii, Arrow) and posterior (Fig.7.A,Aiii, arrowhead) of the embryo. At this stage, we observed weak expression of the ocular/antennal *Ap-hh* stripe (Fig.7.A,Aii, Oc/Ant). From stage 9 onwards *Ap-otd* expression appears as two ventral domains marking the cephalic lobes (Fig,7.B-D, arrow). These *Ap-otd* expression domains overlap with the ocular/antennal *Ap-hh* stripe (Fig.B-Bii) and continue to overlap the ocular stripe following stripe splitting (Fig.7.C-Cii,D-Dii, Oc).

**Figure 7.**
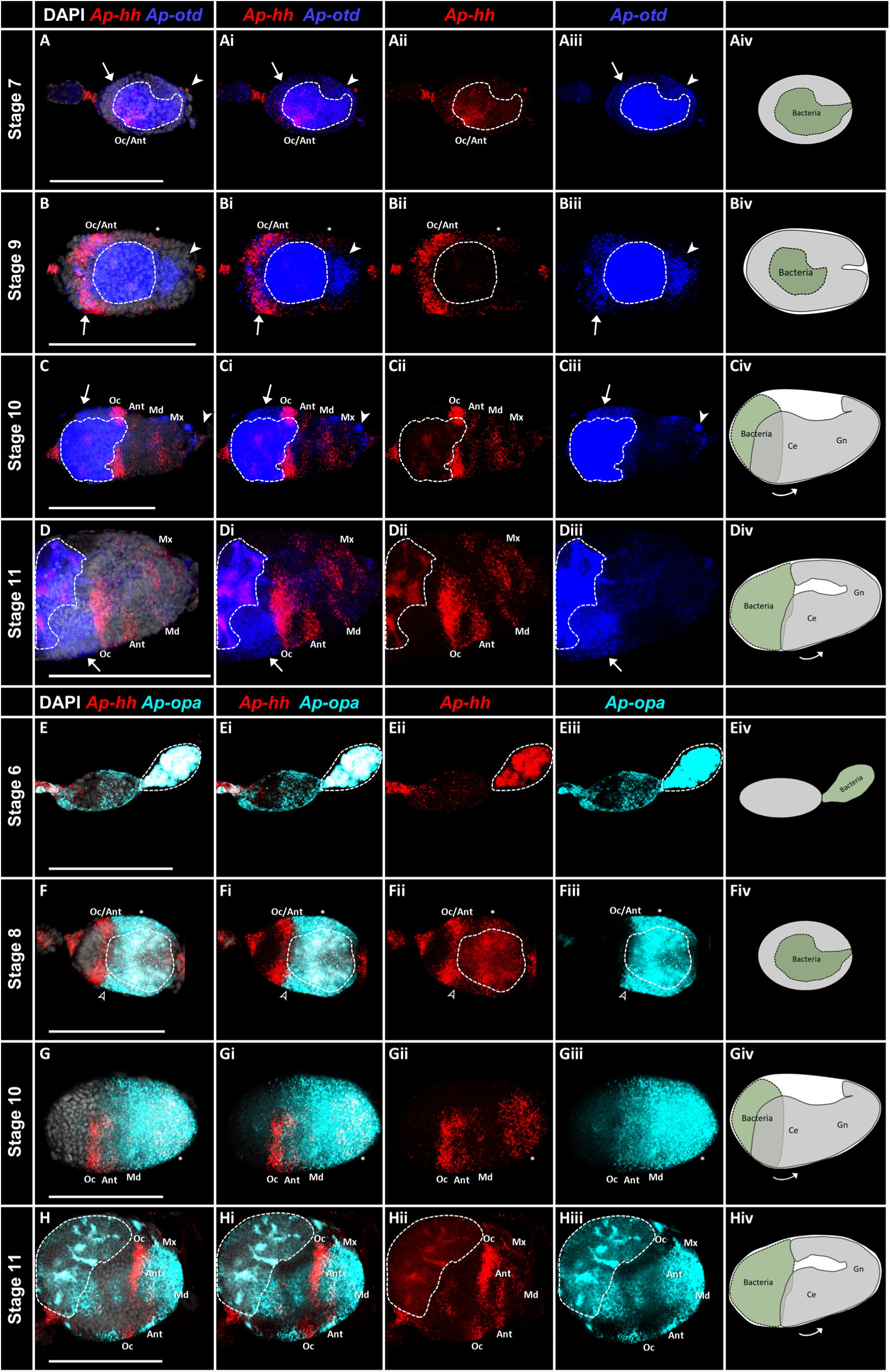
Ocular/antennal *Ap-hh* expression and stripe splitting overlaps expression of *Ap-otd* and *Ap-opa* in viviparous *A. pisum* head segmentation. (A-Hii) Maximum projections of viviparous *A. pisum* embryos of increasing age stained for DAPI (grey), *Ap-hh* (red) and *Ap-otd* (A-Diii, blue) or *Ap-opa* (E-Hiii, cyan). Morphological diagrams (Aiv-Hiv) of corresponding stages are displayed alongside each row to aid interpretation. Each *Ap-hh* stripe is labelled with its corresponding segment: Ocular (Oc), Antennal (Ant) Intercalary (Ic), Mandibular (Md), Maxillary (Mx). All embryos are oriented with the anterior of the embryo to the left. White arrows mark ocular expression of *Ap-otd,* arrowheads mark posterior expression of *Ap-otd.* Black arrowheads mark overlap of Oc/Ant *Ap-hh* stripe and *Ap-opa* expression in stage 8 embryo. White dashed lines mark the position of the bacteriocyte where visible. White asterisk (*) marks posterior *Ap-hh* expression. Scale bars: 100 µm.

Next, we investigated whether *Ap-opa* was expressed alongside the splitting ocular/antennal *Ap-hh* stripe. Firstly, we identified an *A. pisum* homologue of *Tc-opa* (see methods). We then used HCR *in situ* hybridisation to visualise expression of *Ap-hh* and *Ap-opa* throughout head segmentation. *Ap-opa* expression was observed as early as stage 6, preceding the appearance of the ocular/antennal *Ap-hh* stripe, as a broad domain covering the posterior half of the embryo (Fig.7.E,Eiii). By stage 8, *Ap-opa* expression is present as a broad domain from the posterior pole to around 60-70% of embryo length (Fig.7.F,Fiii). The posterior edge of the ocular/antennal *Ap-hh* stripe partially overlaps the *Ap-opa* expression domain at this stage (Fig.7.Fi-Fiii, Oc/Ant, black arrowhead). During stage 10 the splitting *Ap-hh* ocular/antennal stripe overlaps with *Ap-opa* expression (Fig.7.G-Giii, Oc, Ant). By stage 11, *Ap-opa* is expressed overlapping the antennal *Ap-hh* stripe as well as the gap between the ocular and antennal *Ap-hh* stripes (Fig.7.H-Hiii, Oc, Ant). In addition, *Ap-opa* expression appears as two small domains anterior to each *Ap-hh* stripe within the cephalic lobes (Fig.7.H-Hiii, Oc). *Ap-opa* expression also overlaps the mandibular and maxillary *Ap-hh* stripes (Fig.7.H-Hiii, Md, Mx).

Taken together, these results show that *Ap-hh* is expressed partially overlapping an anterior *Ap-otd* expression domain and a posterior *Ap-opa* expression domain throughout *A. pisum* ocular/antennal stripe splitting.

## Discussion

### Conservation of *hh*, *otd* and *opa* expression dynamics during anterior head segmentation in arthropods

Our data reveal remarkable similarities in the expression dynamics of *hh*, *otd* and *opa* homologues in *T. castaneum* and *A. pisum* during procephalon segmentation (Fig.8.A,Ai). In both species, a single ocular/antennal *hh* stripe is expressed partially overlapping an anterior *otd* expression domain and a posterior *opa* expression domain (Fig.2.B-Biiii, Fig.7.B-Bi, F,Fi, Fig.8.A,Ai). This initial *hh* stripe undergoes a single stripe splitting event to produce an anterior ocular *hh* stripe overlapping with *otd* expression and a posterior antennal *hh* stripe (Fig.2.Fii,Fv, Fig.7.Ci,Di, Fig.8.A,Ai). During stripe splitting, *opa* is expressed between the splitting ocular and antennal stripes (Fig.2.Fiv, Fig.7.Gi. Fig.8.A,Ai). These data show that the expression dynamics of each gene are conserved between a holometabolous (*T. castaneum*) and a hemimetabolous (*A. pisum*) insect consistent with conservation across Insecta. The conservation of *hh*, *otd* and *opa* expression during viviparous asexual development in *A. pisum* is particularly striking, given that the segmentation mechanisms operating in the asexual morph otherwise appear evolutionary derived (39).

**Figure 8.**
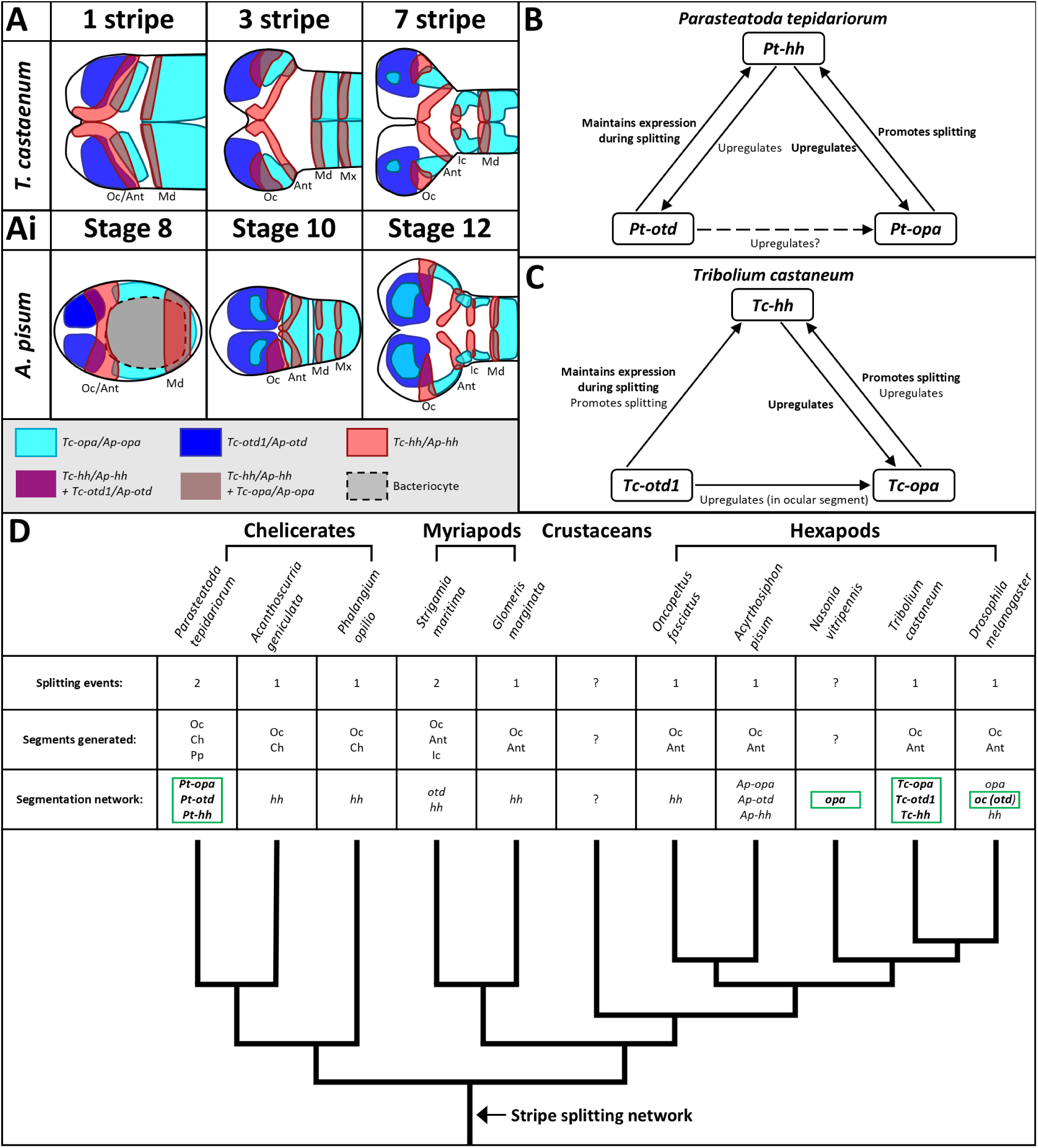
Conservation of the anterior head patterning network across arthropods. (A-Ai) Diagrams depicting the expression domains of *Tc-opa*, *Tc-otd1, Tc-hh* (A) and *Ap-opa*, *Ap-otd*, and *Ap-hh* (Ai) during *T. castaneum* and *A. pisum* head segmentation respectively. (B,C) models of head segmentation network interactions reported for *P. tepidariorum* (B)(Kanayama et al., 2011) and observed in this paper for *T. castaneum* (C). Conserved interactions are in bold. (D) Table summarising the number of *hh* splitting events, the head segments generated by *hh* stripe splitting and evidence for the involvement of *opa*, *otd* and *hh* homologues in head/procephalon development for *P. tepidariorum* (11), *A. geniculata*, *P. opilio* (14), *S. maritima* (12), *G. marginata* (13), *O. fasciatus* (66), *A. pisum* (this study), *N. vitripennis* (19) *T. castaneum* (this study) and *D. melanogaster* (15, 16, 21). Homologues which have been shown to have functional roles in head segmentation are in bold and outlined in green. The phylogeny of each species is displayed with an arrow marking the evolution of the stripe splitting network prior to the divergence of chelicerates and mandibulates.

The positional relationship between *hh*, *otd* and *opa* homologues during stripe splitting in the chelicerate *P. tepidariorum* (11) appears similar to the insect models, with *Pt-hh* expressed overlapping an anterior *Pt-otd* expression domain and a posterior *Pt-opa* expression domain (11). This is consistent with the expression dynamics of *hh*, *otd* and *opa* homologues during head/procephalon segmentation being broadly conserved across arthropods, although studies in other arthropod groups, in particular myriapods and crustaceans, will be required to prove this.

### Conserved roles for *hh*, *otd* and *opa* in anterior head segmentation in arthropods

We analysed the function of *Tc-hh*, *Tc-otd* and *Tc-opa* in *T. castaneum* procephalon segmentation. Similarly to *P. tepidariorum* (11)(Fig.8.B) we found that *Tc-hh* was required for normal *hh* stripe splitting (Fig.5.Hi-Ji, Fig.8.C) and may upregulate expression of *Tc-opa* in the ocular and antennal segments in germband stages (Fig.5. Gii-Jii, Fig.8.C). Furthermore, we found that *Tc-opa* promotes *Tc-hh* stripe splitting (Fig.3.Hi-Ji, Fig.8,C) in a similar manner to *Pt-opa* (Fig.8,B) and *Tc-otd1* maintains expression of *Tc-hh* (Fig.4.F,H, Fig.8.C) following stripe splitting similarly to *Pt-otd* (Fig.8.B) (11).

In contrast to these conserved aspects of head segmentation, we also observed differences in the functional interactions between these genes between *P. tepidarioum* and *T. castaneum*. Firstly, *Tc-hh* is not required for initial expression of *Tc-opa* or *Tc-otd1* which are expressed prior to *Tc-hh* in the *T. castaneum* blastoderm (Fig.2.Ai-Aiii, Fig.8.B.C) (20, 22, 23), which may reflect earlier maternal control of initial *opa* and/or *otd* expression in insects. Secondly, the ocular/antennal *Tc-hh* stripe appeared unsplit in many *Tc-otd* RNAi embryos suggesting that *Tc-otd1* may also promote stripe splitting unlike *Pt-otd* (Fig.4.G, Fig.8.B,C) (11). Furthermore, *Tc-opa* and *Tc-otd1* RNAi embryos displayed phenotypes not observed in *P. terpiadriorum* (11) including reduced expression of *Tc-hh* along the posterior edge of the ocular/antennal stripe (*Tc-*opa, Fig.3.Cii,Dii,O, Fig.8C), reduction of the head lobes (*Tc-opa,* Fig.3.K-M) and deletion of the head field (*Tc-otd1*, Fig.4.I-K).

Similar to *T. castaneum*, a wedge shaped expression domain of *opa* in the anterior blastoderm has been observed in the parasitic wasp *Nasonia vitripennis* (*N. vitripennis*) with RNAi against *Nv-opa* also leading to head reduction phenotypes (19).

Our results, when combined with those of other authors, suggest that *hh*, *otd* and *opa* homologues have conserved roles in Euarthropod head segmentation via regulation of *hh* stripe splitting in both Insecta and Chelicerata, albeit with some species-specific differences in the regulatory interactions between the genes. It will be interesting to see to what extent this patterning gene network is conserved beyond arthropods, given the widely conserved role of these three homologues in anterior and/or head/brain development across animals.

### Implications for our understanding of the origin and evolution of the arthropod head

The splitting of *hh* stripes during the patterning of head/procephalon segments is clearly an ancient and conserved feature of arthropod development (Fig.8.D). Our data strengthen the view that a single *hh* stripe splitting event that patterns the ocular and antennal segments is the ancestral state for insects, with patterning of the developmentally delayed, and morphologically reduced, 3^rd^ procephalic (“intercalary”) segment occurring separately and not via a *hh* stripe splitting mechanism.

Whilst our data don’t help resolve the question of whether one or two *hh* stripe splitting events are ancestral to arthropods (future studies on crustaceans would be particularly informative in that regard), our data do provide evidence that an ancient and conserved gene network involving at least three genes (rather than only the *hh* gene), controls the patterning and formation of the most anterior head segments in arthropods (Fig.8.D). Specifically, overlapping and/or closely abutting expression of Otd/Otx/Oc and Opa appears to be important for driving *hh* stripe splitting and anterior head segmentation. Indeed, a recent expression and genomic study in *Drosophila* suggests that transient overlapping Otd/Otx/Oc and Opa expression likely regulates early head and brain development in fruit flies (21).

Studies similar to this one, on a wider range of arthropods, will be needed to address a number of outstanding questions: For example, has the number of anterior head segments under the control of this ancient and ancestral gene network increased in some lineages (e.g. in *P. tepidariorum* and *S. maritima*)? Or, alternatively, are some segments (e.g. the intercalary segment of insects) no longer patterned via a network that ancestrally patterned the entire head/procephalon? If the latter is true, that might imply the head segments of chelicerates and the procephalic segments of mandibulates originally evolved via serial duplication from one ancestral head segment/region (5, 18).

A third possibility is that a largely conserved head gene network exhibits very different expression dynamics (e.g. stripe splitting vs the *de novo* appearance of single stripes) depending on the nature of upstream regulatory inputs into the network and/or the cellular and temporal/heterochronic context in which segmentation is taking place (see discussion in Janssen (14)), as has already been reported for the trunk segmentation gene network (10, 19, 20, 46). Indeed, it’s curious that the trunk temporal factor Opa appears to be driving frequency doubling (i.e., one stripe of segmentation gene expression becoming two) during both head and trunk segmentation in insects.

### Implications for our understanding of the evolution of anterior-posterior patterning in insects; an evolutionary model

An almost universal feature of animal development is the requirement for Wnt signalling to be repressed in the anterior in order to create a permissive environment for head/brain development (47, 48). It’s perhaps not surprising therefore that Opa, a protein already known to inhibit Wnt signalling and regulate Hedgehog signalling via its role as a transcription factor cofactor (26), is required for *hh* stripe splitting-mediated head segmentation in insects; Opa appears the perfect molecular fit for this role.

In some holometabolous insect species, Wnt signalling pathway components that act to repress Wnt signalling (e.g. Axin in *T. castaneum*; (49), Pangolin in mosquitoes and crane flies; (50), and Panish in a mosquito-like midge; (51)) are maternally localised to the anterior of the egg, presumably acting early and upstream of zygotically expressed Opa to establish a permissive Wnt-free environment for head/brain development (Fig.9.B,C). If one took a holometabolous insect-centric view, one might assume that Opa evolved its zygotic anterior head patterning role after a maternal factor had already evolved to establish Wnt repression in the anterior of the egg. However, a shift from predominantly zygotic patterning to a greater reliance on maternal patterning inputs, and the evolution of an anterior maternal patterning centre, are likely derived features of holometabolous insect development (reviewed in Peel (52); first proposed by Lynch et al. (53)). It’s therefore tempting to speculate that Opa was the ancestral (zygotically expressed) upstream factor that established a Wnt-free, permissive environment for anterior head development (Fig.9.A), with anterior Wnt-repression via maternal factors evolving later in different ways in different holometabolous lineages (Fig. 9.B,C), presumably under selection pressures for faster development.

**Figure 9.**
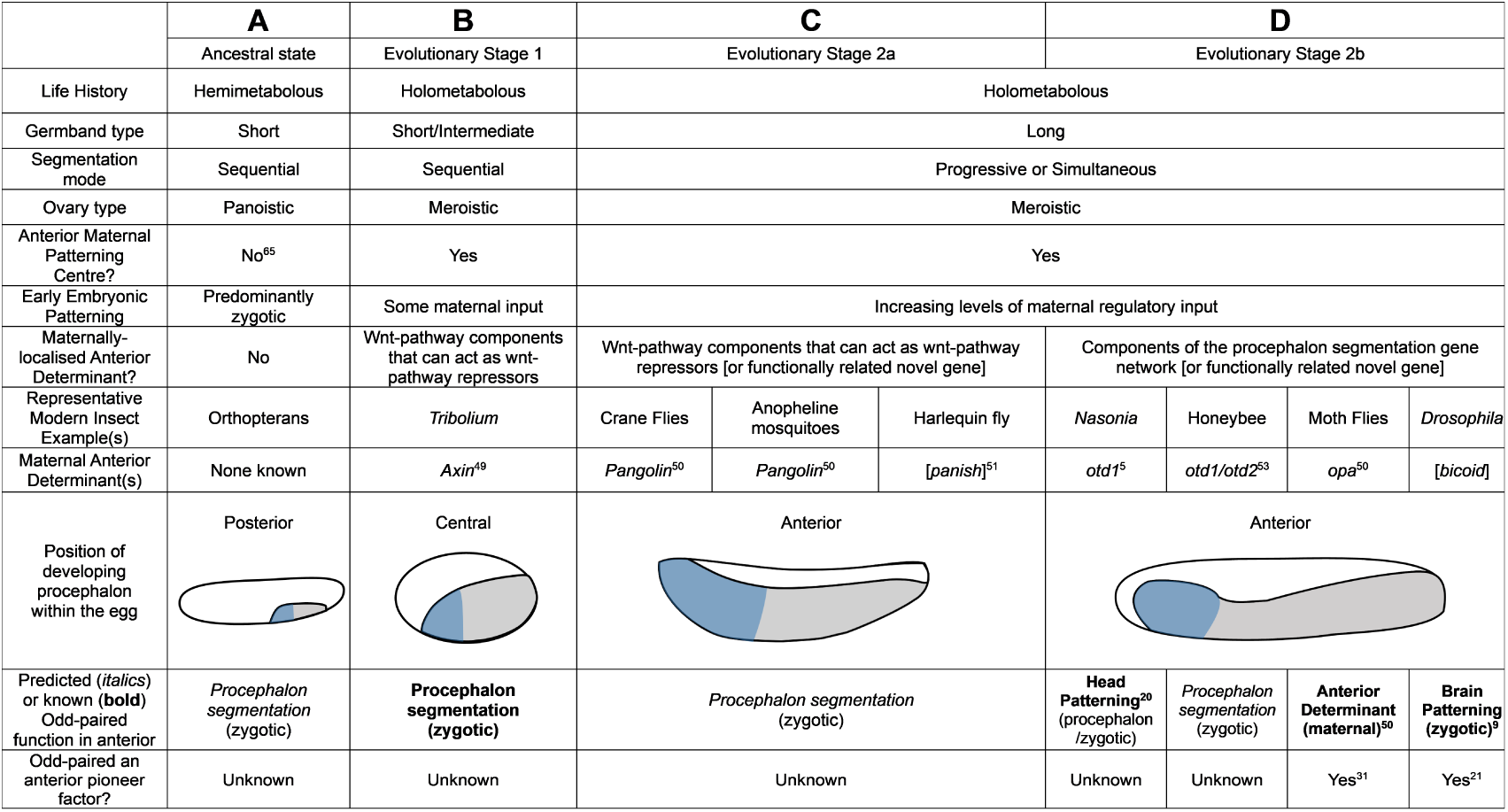
A working model for the evolution of anterior/head patterning in holometabolous insects. (A-D) Comparison of anterior patterning across insects from an assumed ancestral state where the developing procephalon is localised towards the posterior pole of the egg (A) to a more evolutionarily derived central (B, evolutionary stage 1) and then anterior (C, evolutionary stage 2a & D, evolutionary stage 2b) position. Schematic germbands depict the germ rudiment in grey, the developing procephalon in blue and extraembryonic tissue in white. In evolutionary stages 1 (B) and 2a (C), as part of an increase in the maternal patterning of the egg during insect evolution (e.g. the advent of meroistic ovaries), and the transition to long-germ development, Wnt pathway repressors (*axin, pangolin*), or functionally related genes (*panish*), act as maternal anterior determinants, acting upstream of the *odd*/*opa*/*hh* zygotic head patterning network to create a permissive, Wnt-free, environment for head development. In evolutionary stage 2b (D), components of the procephalon segmentation gene network itself (*otd1*/*otd2* and *opa*) are maternally coopted to act as maternal anterior determinants, at least in dipterans in place of wnt repressors, thus shortening and speeding up the regulatory cascade needed for anterior/head patterning. This maternal cooption of components of the procephalon segmentation gene network could only have happened once the developing procephalon had already moved to the anterior pole of the egg and is proposed to have occurred convergently in different long-germ insect lineages. Given the evidence presented in this study for the conservation of the *otd*/*opa*/*hh* head/procephalic patterning gene network between spiders and *Tribolium*, the known role of Opa in Wnt-signalling repression, and the assumption that early patterning in ancestral insects was largely zygotically controlled, we hypothesise that zygotic Opa was ancestrally the primary upstream factor repressing Wnt-signalling in the anterior of the germ rudiment and created a permissive environment for *hh-*driven patterning and development of the anterior head. Whether Opa might have done this by acting as an anterior ‘pioneer factor’ is a future research question. See Discussion for further details. Sources of key information in the table are indicated as superscript citations.

Further to the above, it’s striking that in some ‘long-germ’ holometabolous insect species where the head develops closely abutting the anterior pole of the egg (as opposed to an ancestrally more posterior egg position in ‘short-germ’ insects), it is components of the anterior head patterning network (not Wnt-pathway components) that have been coopted into an early maternal anterior determinant role (Fig.9.D). For example, *opa* is anteriorly localised in moth flies (50), whereas *otd* homologues are maternally localised to the anterior in honeybees (54) and *N. vitripennis* (19, 53). Indeed, Bicoid, the famous *Drosophila* maternal factor, has long been assumed to have substituted for Orthodenticle as an anterior determinant (55–57). Maternal co-option of zygotic anterior head patterning genes might have been an evolutionarily efficient way of speeding up head development in particular, and anterior embryonic development in general, in some long-germ insects (e.g. dipterans), by removing a step in the regulatory cascade (i.e. Wnt-signalling repression by Wnt pathway components). This would explain why a Wnt-signalling gradient is no longer required to set up the anterior-posterior axis in some long-germ insects.

The recent evidence from dipteran species that Odd-paired acts alongside Zelda as an early anterior pioneer factor (not just a transcription factor cofactor) is interesting in the context of this model (21, 31). Whether Odd-paired’s role as an anterior pioneer factor is ancient, and conserved beyond dipterans, or has any relevance to the process of *hh* stripe splitting, remains to be determined (Fig.9D).

## Methods

### T. castaneum methods

#### *T. castaneum* rearing and egg collection

*T. castaneum* beetles (San Bernardino strain) were raised at 30°C on organic wholemeal flour (Doves Farm Foods, Hungerford, UK) with 5% yeast and 0.5% Fumagillin. For egg collection, adult beetles were transplanted onto organic white flour and incubated at 30°C for 24-48 hours. Collected eggs were fixed for *in-situ* Hybridisation Chain Reaction (HCR) following the process detailed by Schinko et al. (58).

### Parental RNAi

*Tc-opa* dsRNA was produced as described previously by Clark and Peel (20). The mRNA sequence of *Tc-hh* and *Tc-otd1* was obtained from iBeetle-base (59). DNA fragments for both genes were amplified by PCR from *T. castaneum* cDNA. For *Tc-hh* the following primer pair was used to produce a 613bp DNA fragment spanning all three exons of the *Tc-hh* gene: 5’-CGCTCGTCTTCAAACAGCATGT-3’ and 5’- TAGAACAAAGTCCGCCGTAGCA-3’. For *Tc-otd1* two DNA fragments were amplified to produce 5’ and 3’ templates for dsRNA. The 5’ fragment was 600bp in length and was produced using the following primer pair: 5’-GCGGCAAGTGTCGGTGAAGGAA-3’ and 5’- GCTCTGCATCTGGGGCGATTCC-3’. The 3’ fragment was 460bp long and was produced using the following primer pair: 5’-CGTTCAACTGGACGGCCAATGG-3’ and 5’- CCATCTGGGTTACAGTCCGCTT-3’. Finally, for *gfp* a single DNA fragment was produced using the following primer pair: 5’-ATGGGTAGTAAAGGAGAAGAACTTT-3’ and 5’-GGGATTACACATGGCATGGA-3’.

DNA fragments from each gene were cloned into the pGEM-Teasy vector (Promega) and used to produce sense and antisense single strand RNA (ssRNA) using the T7 and SP6 MEGAscript High Yield Transcription Kits (Ambion). Sense and antisense ssRNA were then annealed in equimolar amounts and diluted to produce 1 μg/μl stocks of dsRNA for each gene. Stocks were aliquoted and stored at -20°C indefinitely.

Parental RNAi for each gene was carried out using the protocol established by Posnien et al. (60). For *Tc-hh* RNAi, 101 adult females were injected with 5’ *Tc-hh* dsRNA and the same number were injected with *gfp* dsRNA as a control. For *Tc-otd1* RNAi, 117 adult females were injected with 5’ *Tc-otd1* dsRNA, the same number were injected with 3’ *Tc-otd1* dsRNA and 114 were injected with *gfp* dsRNA. For *Tc-opa* RNAi, 119 adult females were injected with 5’ *Tc-opa* dsRNA, 121 adult females were injected with 3’ *Tc-opa* dsRNA and 103 adult females were injected with *gfp* dsRNA. Injected females were left to recover on organic wholemeal flour for one day before being crossed to half their number of adult males on white flour. 24 hours after crossing, eggs were collected from each cross. Each cross was then left on organic wholemeal flour for one day to recover before establishing a new 24 hour egg collection on white flour. This was repeated for a total of eight collections. 10% of the eggs from each collection were allowed to develop and used for hatched and unhatched cuticle analysis. The remaining 90% were fixed as detailed previously.

### Cuticle analysis

Eggs taken for cuticle analysis were left at room temperature for 7-9 days. Any eggs which failed to hatch after one week were dechorionated by immersion in 20% (v/v) bleach, mounted in a 1:1 solution of Hoyer’s medium and lactic acid and incubated at 60°C for up to 48 hours. Cuticles were analysed and imaged using a Leica M165FC Fluorescence Stereo Microscope with a Q Imaging Retiga EXI colour cooled fluorescence camera and Q Capture Pro 7 software.

Unhatched cuticles were analysed for head reduction phenotypes and sorted into one of the following phenotypic categories: wildtype, weak head reduction (loss/reduction of head capsule and or labrum and antennae), strong head reduction (loss/reduction of procephalic and gnathal appendages), reduced head and thorax (loss/reduction of head and thorax segments), wildtype head with abnormal thorax/abdomen and other (cuticle ball and or cuticle fragments).

### Hybridisation chain reaction *in situ* hybridisation

For HCR, template sequences were sourced from NCBI (*Tc-kni,* NM_001128495.1; *Tc-hh,* NM_001114365.1; *Tc-opa,* XM_008203116.2, *Tc- otd1* NM_001039424.1). These sequences were provided to Molecular Instruments which produced fluorescently labelled amplifiers and version 3.0 HCR probes for each gene. The HCR protocol detailed by Bruce et al. (61) was used with the following adjustments: dextran sulphate in the amplification buffer was reduced from 10% (v/v) to 5% (v/v), embryos were incubated with probes for up to 22 hours, the volumes of each probe and amplifier added to embryos were increased to 3µL, embryos were washed for 45 minutes with 2µg/mL DAPI and embryos were stored in a mixture of 2.5% (w/v) DABCO, 50mM tris pH8.0 and 90% glycerol (DTG). Following completion of the HCR protocol, embryos were allowed to rest for one day at 4°C before they were mounted in DTG. Embryos produced from RNAi with morphological phenotypes were selected and mounted along with embryos which displayed wild type morphology.

### Mounting and Imaging

Following HCR, germband embryos were flat-mounted and blastoderm embryos were whole-mounted. Embryos were then imaged using a Zeiss LSM880 + Airyscan Upright Microscope with Zeiss ZEN software at the Faculty of Biological Sciences Bio-imaging and Flow Cytometry facility (University of Leeds). Embryos were imaged as 8-bit Z-stacks using an EC Plan-Neofluor 10x/0.30 objective (no immersion) for whole embryo imaging and a Plan-Apochromat 20x/0.8 objective (no immersion) for imaging of the embryonic head. Z-stacks spanned the entire ventral-dorsal depth of germband embryos and approximately half the depth of blastoderm stage embryos.

Images were analysed and processed using Fiji (62). All Z-stacks were converted into maximum projections. Fiji was also used to rotate, crop and adjust brightness and contrast. Germband embryos were staged based on the number of sequential *Tc-hh* stripes observed in the embryo, counting the mandibular stripe as the first stripe.

### Cell counting

For *Tc-opa* RNAi blastoderm and germband *Tc-hh* analysis, cells were counted using the Fiji cell counter plugin (62). Z-stacks of *Tc-opa* RNAi and *gfp* RNAi embryos stained for DAPI and *Tc-hh*, *Tc-otd1* and *Tc-opa* expression were analysed slice by slice across the entire Z-stack. For blastoderm analysis, three *Tc-opa* RNAi embryos and three *gfp* RNAi embryos of corresponding stages were analysed. For germband analysis, weak *Tc-opa* RNAi embryos which displayed no or minimal head reduction phenotypes were counted. Three germband embryos were analysed for each treatment and each stripe stage, meaning 24 embryos were analysed in total. Cells were counted across the whole procephalon and allocated into four categories: cells within the ocular lobes anterior to the *Tc-hh* stripe(s); cells within the ocular/antennal *Tc-hh* stripe(s), cells in the gap between the splitting ocular/antennal *Tc-hh* stripes; cells between the posterior boundary of the *Tc-hh* stripe and the anterior boundary of mandibular *Tc-hh* and *Tc-opa* expression.

In both blastoderm and *Tc-opa* RNAi analysis, individual cells were marked by DAPI. *Tc-hh* expression was assessed in each cell in the ocular/antennal *Tc-hh* stripe. Cells with at least 30% coverage by *Tc-hh* expression were classified as *Tc-hh* expressing cells in blastoderm and germband stage embryos.

For weak *Tc-opa* RNAi analysis, cell count values were converted into proportional data with each category expressed as a percentage of cells in the total procephalon to correct for differences in embryonic size and reduce the effects of human error. These proportional values were then converted into mean values for each stripe stage. Mean proportional data for *gfp* RNAi and *Tc-opa* RNAi embryos was compared using a paired T-test.

### A. pisum methods

#### *A. pisum* rearing and embryo collection

All aphids were of the N116 strain (63) and were maintained on broad beans (The Sutton variety). Aphids were maintained in a temperature-controlled greenhouse with a long-day photoperiod (16L 21°C : 8D 20°C) at 70% humidity. Plants were regularly replaced and the population was controlled to reduce the production of alate aphids. Viviparous embryos were collected and fixed using the same process described by Duncan et al. (39).

#### Identification of *Ap-opa*

An *A. pisum* homologue of *Tc-opa* was identified through nucleotide BLAST of the *Tc-opa* mRNA sequence (XM_008203116.2) against the pea aphid genome. This identified a predicted ZIC 4 homologue (*Ap-opa,* XM_001943022.5) with a query coverage of 25% and a percent identity of 77.94% (E value: 2e^-91^). Following this, a protein alignment of the *Tc-opa* (XP_008201338) and *Ap-opa* (XP_001943057.1) proteins was performed (Fig.S.3.) which found a query cover of 59% and a percentage identity of 71.62% (E-value: 4e^-115^). Analysis of this aligned region revealed conservation of four Zn finger domains and the putative nucleic acid binding domain. This conserved region displayed only two substitutions between *Tc-opa* and *Ap-opa*, P219-Q251 and V277-I310 with the second substitution being functionally equivalent. Based on the conservation of this functional region, the aligned mRNA sequence (XM_001943022.5) was assigned as the *Tc-opa* homologue *Ap-opa*. A reverse BLAST of the *Ap-opa* mRNA sequence against the *T. castaneum* genome returned *Tc-opa* as the highest scoring match, supporting homology between the two sequences.

#### Hybridisation chain reaction *in situ* hybridisation

Template sequences for each gene were obtained from NCBI and AphidBase (64) (*Ap-Dfd*, XM_001946856.5; *Ap-hh*, XM_001943672.5; *Ap-lab,* XM_029487951.1; *Ap-*opa, XM_001943022.5; Ap*-otd*, GQ144357.1) and supplied to Molecular Instruments to produce version 3.0 probes and fluorescently tagged amplifiers.

*A. pisum* embryos underwent HCR following an adaption of the protocol detailed by Bruce et al. (61) with the following changes. After the PTw washes in step 1, embryos were re-fixed in 4% formaldehyde for 20 minutes and washed three times for 10 minutes in PTw. Embryos were permeabilised in detergent solution for 45 and subsequently underwent seven 5 minute washes in PTw before pre-hybridisation. Embryos were incubated with probes for up to 22 and the volumes of each probe and amplifier were increased to 3 µL.

### Mounting and imaging

Following HCR, *A. pisum* embryos were staged according to the stages described by Miura et al. (38). Embryos were flat-mounted in DTG and imaged as described for *T.castaneum* embryos. Z-stack images were captured for embryos covering the entire depth of the embryo. Images were converted into maximum projections and brightness and contrast were adjusted using Fiji (62). Fiji was also used to count the number of cells in the ocular/antennal *Ap-hh* stripe. Individual cells were labelled with DAPI. Cells with at least 30% coverage by *Ap-hh* expression were classified as *Ap-hh* expressing cells.

## Bibliography

1. Eriksson BJ, Tait NN, Budd GE, Janssen R, Akam M. Head patterning and Hox gene expression in an onychophoran and its implications for the arthropod head problem. Development genes and evolution. 2010;220:117–22.

2. Budd GE. A palaeontological solution to the arthropod head problem. Nature. 2002;417(6886):271–5.

3. Rempel J. Evolution of the insect head: the endless dispute. Quaestiones Entomologicae. 1975.

4. Hughes CL, Kaufman TC. Exploring the myriapod body plan: expression patterns of the ten Hox genes in a centipede. Development. 2002;129(5):1225–38.

5. Lev O, Edgecombe GD, Chipman AD. Serial homology and segment identity in the arthropod head. Integrative Organismal Biology. 2022;4(1):obac015.

6. Gainett G, Klementz BC, Blaszczyk PO, Bruce HS, Patel NH, Sharma PP. Dual Functions of labial Resolve the Hox Logic of Chelicerate Head Segments. Molecular Biology and Evolution. 2023;40(3).

7. Telford MJ, Thomas RH. Expression of homeobox genes shows chelicerate arthropods retain their deutocerebral segment. Proceedings of the National Academy of Sciences. 1998;95(18):10671–5.

8. Mittmann B, Scholtz G. Development of the nervous system in the “head” of Limulus polyphemus (Chelicerata: Xiphosura): morphological evidence for a correspondence between the segments of the chelicerae and of the (first) antennae of Mandibulata. Dev Genes Evol. 2003;213(1):9–17.

9. Scholtz G, Edgecombe GD. The evolution of arthropod heads: reconciling morphological, developmental and palaeontological evidence. Development Genes and Evolution. 2006;216(7):395–415.

10. Clark E, Peel AD, Akam M. Arthropod segmentation. Development. 2019;146(18):dev170480.

11. Kanayama M, Akiyama-Oda Y, Nishimura O, Tarui H, Agata K, Oda H. Travelling and splitting of a wave of hedgehog expression involved in spider-head segmentation. Nature Communications. 2011;2:500.

12. Hunnekuhl VS, Akam M. Formation and subdivision of the head field in the centipede Strigamia maritima, as revealed by the expression of head gap gene orthologues and hedgehog dynamics. Evodevo. 2017;8:18.

13. Janssen R. Segment polarity gene expression in a myriapod reveals conserved and diverged aspects of early head patterning in arthropods. Developmental genes and evolution. 2012;222(5):299–309.

14. Janssen R. Early expression of chelicerate hedgehog orthologs and its bearing on the homology of arthropod head segments. Discover Developmental Biology. 2025;235(1):1–10.

15. Lee JJ, von Kessler DP, Parks S, Beachy PA. Secretion and localized transcription suggest a role in positional signaling for products of the segmentation gene hedgehog. Cell. 1992;71(1):33–50.

16. Ntini E, Wimmer EA. Unique establishment of procephalic head segments is supported by the identification of cis-regulatory elements driving segment-specific segment polarity gene expression in Drosophila. Development genes and evolution. 2011;221(1):1–16.

17. Telford MJ, Thomas RH. Demise of the Atelocerata? Nature. 1995;376(6536).

18. Ortega-Hernández J, Janssen R, Budd GE. Origin and evolution of the panarthropod head–a palaeobiological and developmental perspective. Arthropod structure & development. 2017;46(3):354–79.

19. Taylor SE, Dearden PK. The Nasonia pair-rule gene regulatory network retains its function over 300 million years of evolution. Development. 2022;149(5):dev199632.

20. Clark E, Peel AD. Evidence for the temporal regulation of insect segmentation by a conserved sequence of transcription factors. Development. 2018;145(10).

21. Fenelon KD, Gao F, Borad P, Abbasi S, Pachter L, Koromila T. Cell-specific occupancy dynamics between the pioneer-like factor Opa/ZIC and Ocelliless/OTX regulate early head development in embryos. Frontiers in Cell and Developmental Biology. 2023;11:1126507.

22. Schinko JB, Kreuzer N, Offen N, Posnien N, Wimmer EA, Bucher G. Divergent functions of orthodenticle, empty spiracles and buttonhead in early head patterning of the beetle Tribolium castaneum (Coleoptera). Developmental Biology. 2008;317(2):600–13.

23. Kotkamp K, Klingler M, Schoppmeier M. Apparent role of Tribolium orthodenticle in anteroposterior blastoderm patterning largely reflects novel functions in dorsoventral axis formation and cell survival. Development. 2010;137(11):1853–62.

24. Sen S, Reichert H, VijayRaghavan K. Conserved roles of ems/Emx and otd/Otx genes in olfactory and visual system development in Drosophila and mouse. Open Biology. 2013;3(5):120177.

25. Simeone A, Puelles E, Acampora D. The Otx family. Curr Opin Genet Dev. 2002;12(4):409–15.

26. Houtmeyers R, Souopgui J, Tejpar S, Arkell R. The ZIC gene family encodes multi-functional proteins essential for patterning and morphogenesis. Cell Mol Life Sci. 2013;70(20):3791–811.

27. Koromila T, Gao F, Iwasaki Y, He P, Pachter L, Gergen JP, et al. Odd-paired is a pioneer-like factor that coordinates with Zelda to control gene expression in embryos. Elife. 2020;9.

28. Soluri IV, Zumerling LM, Payan Parra OA, Clark EG, Blythe SA. Zygotic pioneer factor activity of Odd-paired/Zic is necessary for late function of the Drosophila segmentation network. Elife. 2020;9.

29. Clark E, Battistara M, Benton MA. A timer gene network is spatially regulated by the terminal system in the Drosophila embryo. Elife. 2022;11:e78902.

30. Clark E, Akam M. Odd-paired controls frequency doubling in Drosophila segmentation by altering the pair-rule gene regulatory network. Elife. 2016;5:e18215.

31. Amiri EE, Tenger-Trolander A, Li M, Julian AT, Kasan K, Sanders SA, et al. Breaking anterior-posterior symmetry in the moth fly *Clogmia albipunctata*. bioRxiv. 2025:2025.01.13.632851.

32. Posnien N, Schinko JB, Kittelmann S, Bucher G. Genetics, development and composition of the insect head--a beetle’s view. Arthropod structure & development. 2010;39(6):399–410.

33. Posnien N, Koniszewski ND, Hein HJ, Bucher G. Candidate gene screen in the red flour beetle Tribolium reveals six3 as ancient regulator of anterior median head and central complex development. PLoS Genet. 2011;7(12):e1002416.

34. Coulcher JF, Telford MJ. Cap’n’collar differentiates the mandible from the maxilla in the beetle Tribolium castaneum. EvoDevo. 2012;3(1):25.

35. Farzana L, Brown SJ. Hedgehog signaling pathway function conserved in Tribolium segmentation. Developmental genes and evolution. 2008;218(3-4):181–92.

36. Li Y, Brown SJ, Hausdorf B, Tautz D, Denell RE, Finkelstein R. Two orthodenticle-related genes in the short-germ beetle Tribolium castaneum. Development genes and evolution. 1996;206:35–45.

37. Kollmann M, Minoli S, Bonhomme J, Homberg U, Schachtner J, Tagu D, et al. Revisiting the anatomy of the central nervous system of a hemimetabolous model insect species: the pea aphid Acyrthosiphon pisum. Cell and tissue research. 2011;343:343–55.

38. Miura T, Braendle C, Shingleton A, Sisk G, Kambhampati S, Stern DL. A comparison of parthenogenetic and sexual embryogenesis of the pea aphid Acyrthosiphon pisum (Hemiptera: Aphidoidea). Journal of Experimental Zoology Part B: Molecular and Developmental Evolution. 2003;295(1):59–81.

39. Duncan EJ, Leask MP, Dearden PK. The pea aphid (Acyrthosiphon pisum) genome encodes two divergent early developmental programs. Developmental Biology. 2013;377(1):262–74.

40. Huang TY, Cook CE, Davis GK, Shigenobu S, Chen RP, Chang CC. Anterior development in the parthenogenetic and viviparous form of the pea aphid, Acyrthosiphon pisum: hunchback and orthodenticle expression. Insect molecular biology. 2010;19 Suppl 2:75–85.

41. Peel AD, Schanda J, Grossmann D, Ruge F, Oberhofer G, Gilles AF, et al. Tc-knirps plays different roles in the specification of antennal and mandibular parasegment boundaries and is regulated by a pair-rule gene in the beetle Tribolium castaneum. BMC Developmental Biology. 2013;13:25.

42. Rogers BT, Peterson MD, Kaufman TC. The development and evolution of insect mouthparts as revealed by the expression patterns of gnathocephalic genes. Evolution & development. 2002;4(2):96–110.

43. Brown SJ, DeCamillis M, Gonzalez-Charneco K, Denell M, Beeman R, Nie W, et al. Implications of the Tribolium *Deformed* mutant phenotype for the evolution of Hox gene function. Proceedings of the National Academy of Sciences. 2000;97(9):4510–4.

44. Posnien N, Bucher G. Formation of the insect head involves lateral contribution of the intercalary segment, which depends on Tc-labial function. Developmental Biology. 2010;338(1):107–16.

45. Economou AD, Telford MJ. Comparative gene expression in the heads of Drosophila melanogaster and Tribolium castaneum and the segmental affinity of the Drosophila hypopharyngeal lobes. Evolution & development. 2009;11(1):88–96.

46. Cheatle Jarvela AM, Trelstad CS, Pick L. Anterior-posterior patterning of segments in Anopheles stephensi offers insights into the transition from sequential to simultaneous segmentation in holometabolous insects. J Exp Zool B Mol Dev Evol. 2023;340(2):116–30.

47. Petersen CP, Reddien PW. Wnt signaling and the polarity of the primary body axis. Cell. 2009;139(6):1056–68.

48. Niehrs C. On growth and form: a Cartesian coordinate system of Wnt and BMP signaling specifies bilaterian body axes. Development. 2010;137(6):845–57.

49. Ansari S, Troelenberg N, Dao VA, Richter T, Bucher G, Klingler M. Double abdomen in a short-germ insect: Zygotic control of axis formation revealed in the beetle Tribolium castaneum. Proceedings of the National Academy of Sciences. 2018;115(8):1819–24.

50. Yoon Y, Klomp J, Martin-Martin I, Criscione F, Calvo E, Ribeiro J, et al. Embryo polarity in moth flies and mosquitoes relies on distinct old genes with localized transcript isoforms. Elife. 2019;8:e46711.

51. Klomp J, Athy D, Kwan CW, Bloch NI, Sandmann T, Lemke S, et al. A cysteine-clamp gene drives embryo polarity in the midge Chironomus. Science. 2015;348(6238):1040–2.

52. Peel AD. The evolution of developmental gene networks: lessons from comparative studies on holometabolous insects. Philosophical Transactions of the Royal Society B: Biological Sciences. 2008;363(1496):1539–47.

53. Lynch JA, Brent AE, Leaf DS, Anne Pultz M, Desplan C. Localized maternal orthodenticle patterns anterior and posterior in the long germ wasp Nasonia. Nature. 2006;439(7077):728–32.

54. Wilson MJ, Dearden PK. Diversity in insect axis formation: two orthodenticle genes and hunchback act in anterior patterning and influence dorsoventral organization in the honeybee (Apis mellifera). Development. 2011;138(16):3497–507.

55. Schröder R. The genes orthodenticle and hunchback substitute for bicoid in the beetle Tribolium. Nature. 2003;422(6932):621–5.

56. Lynch J, Desplan C. Evolution of development: beyond bicoid. Current Biology. 2003;13(14):R557–R9.

57. Treisman J, Gonczy P, Vashishtha M, Harris E, Desplan C. A single amino acid can determine the DNA binding specificity of homeodomain proteins. Cell. 1989;59(3):553–62.

58. Schinko JB, Posnien N, Kittelmann S, Koniszewski N, Bucher G. Single and double whole-mount in situ hybridization in red flour beetle (Tribolium) embryos. Cold Spring Harbor Protocols. 2009;2009(8):pdb. prot5258.

59. Dönitz J, Schmitt-Engel C, Grossmann D, Gerischer L, Tech M, Schoppmeier M, et al. iBeetle-Base: a database for RNAi phenotypes in the red flour beetle Tribolium castaneum. Nucleic acids research. 2015;43(D1):D720–D5.

60. Posnien N, Bashasab F, Bucher G. The insect upper lip (labrum) is a nonsegmental appendage-like structure. Evolution & development. 2009;11(5):480–8.

61. Bruce HS, Jerz G, Kelly S, McCarthy J, Pomerantz A, Senevirathne G, et al. Hybridization chain reaction (HCR) in situ protocol. protocols io. 2021.

62. Schindelin J, Arganda-Carreras I, Frise E, Kaynig V, Longair M, Pietzsch T, et al. Fiji: an open-source platform for biological-image analysis. Nature methods. 2012;9(7):676–82.

63. Kanvil S, Powell G, Turnbull C. Pea aphid biotype performance on diverse Medicago host genotypes indicates highly specific virulence and resistance functions. Bulletin of entomological research. 2014;104(6):689–701.

64. Legeai F, Shigenobu S, Gauthier JP, Colbourne J, Rispe C, Collin O, et al. AphidBase: a centralized bioinformatic resource for annotation of the pea aphid genome. Insect molecular biology. 2010;19:5–12.

65. Sander K. Specification of the basic body pattern in insect embryogenesis. Advances in insect physiology. 1976;12:125–238.

66. Lev O, Chipman AD. Development of the Pre-gnathal Segments in the Milkweed Bug Oncopeltus fasciatus Suggests They Are Not Serial Homologs of Trunk Segments. Frontiers in Cell and Developmental Biology. 2021;9:695135.

