## Supplementary figures for "Getting a head: Evidence for a conserved anterior head patterning gene network in arthropods"

### Supplementary material

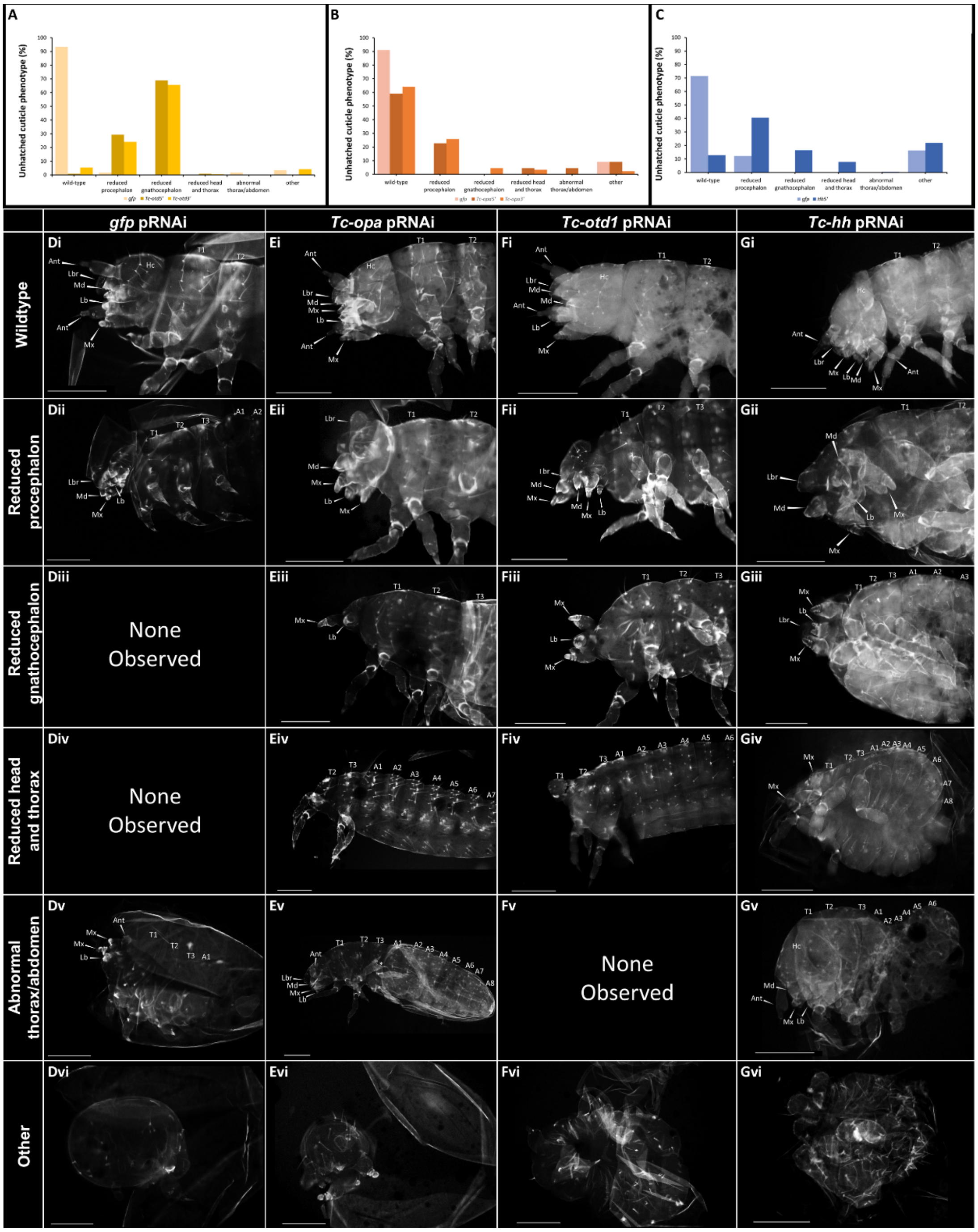


**Figure S.1. Parental RNAi against *Tc-otd1*, *Tc-opa* and *Tc-hh* caused head reduction phenotypes.** (A-C) Percentage of each phenotype observed in *Tc-otd1*, *Tc-opa* and *Tc-hh* RNAi cuticles compared against *gfp* RNAi cuticles. (D-Gvi). Representative unhatched cuticles for each phenotype observed in each condition. Each segment is labelled as follows: Ocular (Oc), Antennal (Ant) Intercalary (Ic), Mandibular (Md), Maxillary (Mx) and Labial (Lb), Thoracic 1-3 (T1-T3), Abdominal 1-8 (A1-A8). Scale bars: 100 µm.

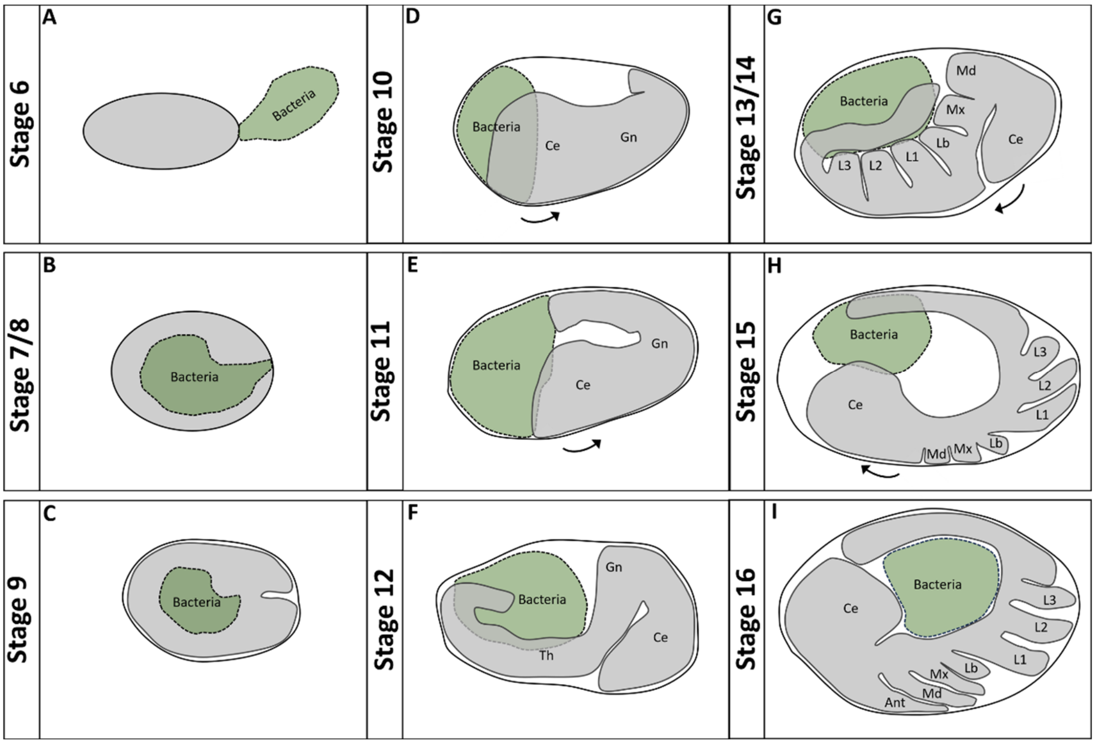


**Figure S.2. Diagram of asexual viviparous *A. pisum* embryogenesis.** (A-I) Simplified diagrams of asexual viviparous *A. pisum* embryogenesis from stage 6 (A) to stage 16 (I). Embryonic tissue is coloured grey whilst the bacteriocyte is coloured green and labelled. At stage 6, the bacteriocyte is present attached to the posterior of the embryo (A). At stage 7, the bacteriocyte is incorporated into the embryo (B). At stage 9, the embryo invaginates at the posterior pole (C). At stage 10, the germband forms and begins extending through stage 11 onwards (D,E). Between stages 10 and 11, the cephalic lobes shift towards the posterior pole (D,E). As the limbs form from stages 13-16, the cephalic lobes bend and shift towards the anterior pole (F-I). Arrows indicate the movements of the cephalic lobes throughout embryogenesis. Approximate regions of the embryo are labelled: (Ce) cephalic lobe, (Gn) gnathal segments, (Th) thoracic segments, (Ant) antenna, (Md) mandible, (Mx) maxilla, (Lb) labium, (L1/2/3) leg 1/2/3. Anterior is left, posterior is right.

| Score | Expect | Method | Identities | Positives | Gaps |
|---|---|---|---|---|---|
| 331 bits(848) | 4e-115 | Compositional matrix adjust. | 164/229(72%) | 180/229(78%) | 7/229(3%) |

```
Tc-opa  113  NPMGVPAHHSHGAFFRYMRQ-PIK----QEMQCLWVDPEQPPPRKICAKHFTSMHEIVTH  167
             +P G     H  GAF+RY R  P+      ++M CLW+D    PP R  C K F SM +IV+H
Ap-opa  141  DPYGCGGGHPAGAFYRYTRHGPVAAATLKDMSCLWIDKPGPPMR-TCGKMFGSMQDIVSH  199

Tc-opa  168  LTVEHVGGPECTTHACFWQNCSRNGRPFKAKYKLVNHIRVHTGEKPFPCPFPGCGKVFAR  227
             +TVEHVGGPECTTHACFW   C R GRPFKAKYKLVNHIRVHTGEKPFPCPF GCGKVFAR
Ap-opa  200  ITVEHVGGPECTTHACFWHGCERKGRPFKAKYKLVNHIRVHTGEKPFPCPFQGCGKVFAR  259

Tc-opa  228  SENLKIHKRTHTGEKPFKCEYEGCDRRFANSSDRKKHSHVHTSDKPYNCRVSGCDKSYTH  287
             SENLKIHKRTHTGEKPFKCEYEGCDRRFANSSDRKKHSHVHTSDKPYNCR+SGCDKSYTH
Ap-opa  260  SENLKIHKRTHTGEKPFKCEYEGCDRRFANSSDRKKHSHVHTSDKPYNCRISGCDKSYTH  319

Tc-opa  288  PSSLRKHMKVHGCSGRSPPHYDSDGEESNSSSAGSISVAASPQVSGPQV   336
             PSSLRKHMKVHG +G+       + D E+SN SS GSI V  S   + P +
Ap-opa  320  PSSLRKHMKVHG-NGKMDSDGNYDSEDSNCSSGGSIRVTDSCTTATPSI   367
```

C2H2 Zn finger domain   Putative nucleic acid binding domain

**Figure.S.3. Alignment of *Tc-opa* and *Ap-opa* protein sequences.** Alignment performed via BLASTP. Predicted C2H2 Zn finger domains are highlighted in blue, the putative nucleic acid binding domain is highlighted in red.
